# PCIF1 is partly cytoplasmic, dynamically localizes to stress granules and binds mRNA coding regions upon oxidative stress

**DOI:** 10.1101/2024.05.08.593175

**Authors:** Trinh T. Tat, Sabeen Raza, Shaheerah Khan, Tiara L. Watson, Sung Yun Jung, Daniel L. Kiss

## Abstract

PCIF1 (*P*hosphorylated *C*TD-*I*nteracting *F*actor *1*) is the mRNA (2’-O-methyladenosine-N(6)-)-methyltransferase that catalyzes the formation of cap-adjacent N_6_,2’-O-dimethyladenosine (m6Am) by methylating adenosines at the first transcribed position of capped mRNAs. While previous studies assumed that PCIF1 was nuclear, cell fractionation and immunofluorescence both show that a population of PCIF1 is localized to the cytoplasm. Further, PCIF1 redistributes to stress granules upon oxidative stress. Immunoprecipitation studies with stressed cells show that PCIF1 also physically interacts with G3BP and other stress granule components. In addition, PCIF1 behaves as a stress granule component as it disassociates from stress granules upon recovery from stress. Overexpressing full-length PCIF1 also inhibits stress granule formation, while knocking out PCIF1 slows stress granule disassembly. Next, our enhanced crosslinking and immunoprecipitation (eCLIP) data show that PCIF1 binds mRNAs in their coding sequences rather than cap-proximal regions. Further PCIF1’s association with mRNAs increased upon NaAsO_2_ stress. In contrast to eCLIP data, ChIP-Seq experiments show that PCIF1 is predominantly associated with transcription start sites rather than gene bodies, indicating that PCIF1’s association with mature mRNA is not co-transcriptional. Collectively, our data suggest that PCIF1 has cytoplasmic RNA surveillance role(s) independent of transcription-associated cap-adjacent mRNA modification, particularly during the stress response.

---

Comparative transcriptomics has revealed the complexity of RNA with a variety of post-transcriptional modifications involving the addition and/or removal of unique chemical moieties^1^. Methylation, or the addition of a methyl group to distinct positions on the nucleobase and/or its ribose sugar moiety, is one of the most varied and ubiquitous RNA modifications^2^. Methylation events often convey specific information guiding the fate of the RNA, including its stability and translation^2, 3^. Transcriptome-wide approaches have shown that epitranscriptomic marks like N6-methyladenosine (m^6^A), 5-methylcytidine (m^5^C), N1-methyladenine (m^1^A), pseudouridine, are both widespread on mRNAs and confer at least some manner of regulation to their host mRNAs^1^.

Although first identified in mRNAs nearly five decades ago, N6, 2’-O-dimethyladenosine (m^6^Am) has mostly escaped attention until recently^4^. Unlike m^6^A, which is generally deposited along mRNA body with peak modification density observed near the stop codon, m^6^Am marks were found at the first nucleotide after the 5’-N7-methylguanosine (m^7^G) cap of mRNAs^4, 5^. The in vivo role(s) of m^6^Am remains a matter of debate as recent studies have generated opposing conclusions^3, 6^. First, mRNAs with cap-adjacent m^6^Am modifications were more resistant to decapping by Dcp2, consistent with m^6^Am conferring increased mRNA stability^3^. In contrast, more recent works have shown that the presence of m^6^Am had either little or a negative effect on stability of mRNAs, and the gene expression was also m^6^Am independent^6–8^. The likeliest interpretation of these conflicting data is that the function of m^6^Am on mRNAs doesn’t follow a single rule. Rather, m^6^Am’s influence on mRNA stability and/or translation is likely context dependent.

Shortly after m^6^Am was identified, cell fractionation studies were able to isolate the biochemical activity responsible for adding cap-adjacent m^6^Am, however the technologies of the day prevented the identification of the responsible enzyme^9^. This changed several years ago when several teams identified Phosphorylated CTD Interacting Factor 1 (PCIF1) as the writer of m^6^Am modifications^6, 7, 10, 11^. PCIF1 was first identified in 2003 as associating with the C-terminal domain of RNA Polymerase II^12, 13^. Cap-adjacent m^6^Am is formed when PCIF1 uses S-adenosylmethionine to catalyze the N6-methylation of cap-adjacent 2’-O-methyladenosine^6^. PCIF1-knockout cells have been used to identify the location and functions of m6Am marks in the transcriptome^7, 10^. Further, using PCIF1-depleted cells or m^6^Am-Exo-Seq which leverages the exonucleolytic susceptibility of uncapped RNAs, mapped the distribution of m^6^A and m^6^Am marks across the transcriptome^7, 14^. In addition, PCIF1^-/-^ mice revealed that mammalian PCIF1 regulated transcript levels in mouse tissues, including testes, brain, and spleen^15^. Furthermore, PCIF1 and/or m^6^Am have been implicated in prostate and other cancers^16, 17^. Depending upon the virus, PCIF1 can aid or hinder viral infection^14, 18^. As we have been interested in stress responses for some time, found these links to viral infections, a type of cellular stressor, particularly interesting.

The integrated stress response (ISR) is a complex and coordinated reaction to different external stressors including oxidative and proteotoxic stresses among others^19^. The different triggers of the ISR funnel through one of four eukaryotic translation initiation factor 2 α (eIF2α) kinases which causes a global shutdown in translation^19^. In addition to translation shutdown, the formation of stress granules, dynamic, membrane-less organelles composed of protein and RNA aggregates are hallmarks of the stress response^19, 20^. Stress granules are thought to play a role in modulating RNA metabolism, mRNA translation, and cellular homeostasis during stress^21^. Stress granule assembly plays important roles in shifting cellular translation to stress response mRNAs and shut off globally cap-dependent translation^20, 22^. Recent works have revealed that stress granules have both core and stress-specific components, and that stress granules contain mostly translationally silenced mRNAs^22, 23^.

While the functions of m^6^Am on mRNA processing remain debated, our study examined PCIF1, the primary writer of m^6^Am, and whether it has a role(s) in post-transcriptional gene regulation in the cytoplasm. Particularly perplexing were the methods needed to isolate and track PCIF1’s activity in vitro. The works that identified PCIF1 as the cells’ m^6^Am writer used either cytoplasmic^9, 11^ or whole cell extracts^6, 10^, not nuclear extracts as expected for a co-transcriptional process. We reasoned that PCIF1 could have distinct sub-cellular compartment-specific activities in vivo, and that some of these activities would be cytoplasmic. Our data here confirm our earlier observation that a fraction of PCIF1 is always cytoplasmic^24^. We show that PCIF1 both localizes to stress granules formed by oxidative stress and interacts with a significant number of stress granule resident proteins. We also demonstrate that PCIF1 directly binds stress granule resident mRNAs, and that the number of mRNAs bound increases markedly upon oxidative stress. Further, in contrast to prior observations based on the location of m^6^Am marks, our data show that PCIF1 is primarily bound to coding regions, not only to 5’ untranslated regions (UTRs).

## Results

### PCIF1 is both nuclear and cytoplasmic in mammalian cells

PCIF1 was originally identified and named for its interactions with the phosphorylated C-terminal domain (CTD) of RNA polymerase II via its WW domain^12, 13^. Therefore, despite readily available contradictory evidence from the human protein atlas and other sources, it was assumed to localize in the nucleus and be limited to roles in nuclear mRNA modification^25^. We used both cell fractionation experiments and indirect immunofluorescence (IF) to definitively ascertain the distribution of PCIF1 in different cell lines. Interestingly, our data reveal that PCIF1 is present in the cytoplasm in HUVEC, U2OS, and HEK293T cell lines (Fig. 1A). We next used a combination of immunofluorescence staining and confocal microscopy to confirm these observations. Our resulting data from HUVEC and HEK293 cell lines (Fig. 1B) show that a population of PCIF1 is indeed cytoplasmic, that our antibodies have no non-specific signal (Fig S1), and are consistent with updated classifications in the human protein atlas^25^. We were vigilant about not basing our conclusions on a single antibody, so we validated their specificity using PCIF1 knockouts (Fig 1A and S2) and tested several commercially available PCIF1 antibodies (Fig. S2, S5 and Table S1) for both Western Blotting and IF. Interestingly, we also found that PCIF1 co-localized with PDI (Protein disulfide isomerase), which is a commonly used endoplasmic reticulum marker, in both HUVEC and HEK293 cells (Fig. S3).

**Fig 1.**
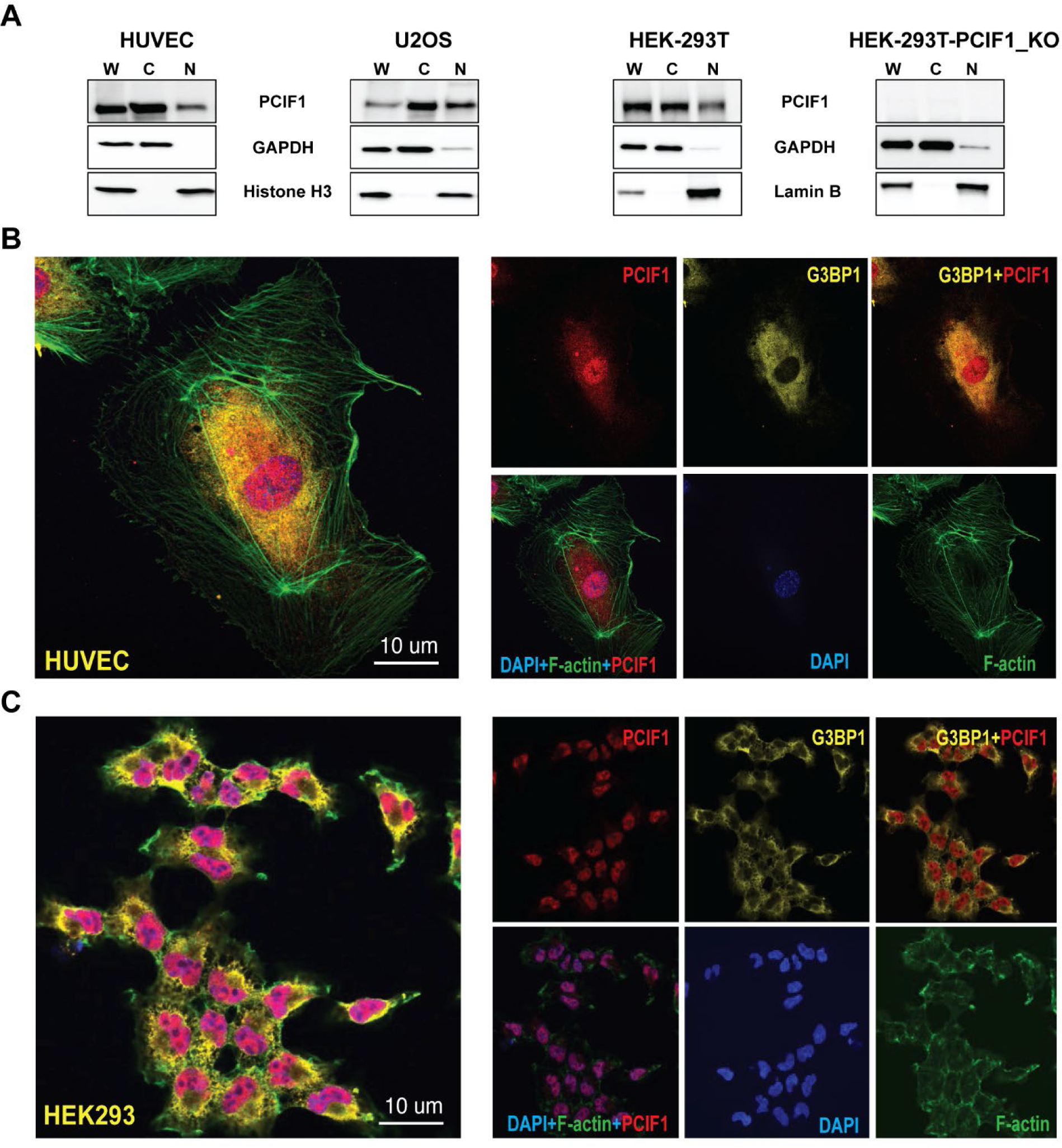
PCIF1 is predominantly found in cytoplasmic compartment in a variety of mammalian cell types. (A) Cells were fractionated and whole cell (W), cytoplasmic (C) and nuclear (N) extracts from the indicated cell lines were assayed by Western Blotting with antibodies targeting PCIF1 and cytoplasmic (GAPDH) or nuclear (Histone H3 or Lamin B) markers. (B) HUVEC and (C) HEK293 cells were grown on coverslips and stained with antibodies targeting PCIF1 (red), G3BP1 (yellow), phalloidin (green) and DAPI (blue). Images represent the consensus data from three independent biological replicates.

### PCIF1 dynamically redistributes to stress granules during stress and disassociates from stress granules when the cells recover from stress

As mentioned above, PCIF1 has been linked to viral infections^14, 18^. Since a portion of PCIF1 was cytoplasmic, we tested if it redistributed during the stress response. HUVEC and HEK293 cells were treated with sodium arsenite (NaAsO_2_) at 500 µM for up to 120 minutes. Translation shutdown was observed using polysome gradients and (Fig S4) the appearance of G3BP1-containing foci validate that our conditions successfully induced a stress response. The resulting data (Fig 2, S4, S5) clearly show that PCIF1 (labeled in red) concentrates in distinct cytoplasmic foci that fully overlap with foci observed by targeting the canonical stress granule protein G3BP1 (Ras GTPase-activating protein binding, yellow) during the NaAsO_2_ induced stress response. Both HUVEC and HEK293 cells with NaAsO_2_ for 1 hour show nearly identical localization patterns (Figure 2A). Notably, the same staining pattern was reproduced using four different PCIF1-targeting antibodies directed against different regions of PCIF1 (Fig S5, Table S1). This co-residence in stress granules could portend a potential role for PCIF1 in stress granule formation and/or function.

**Fig 2.**
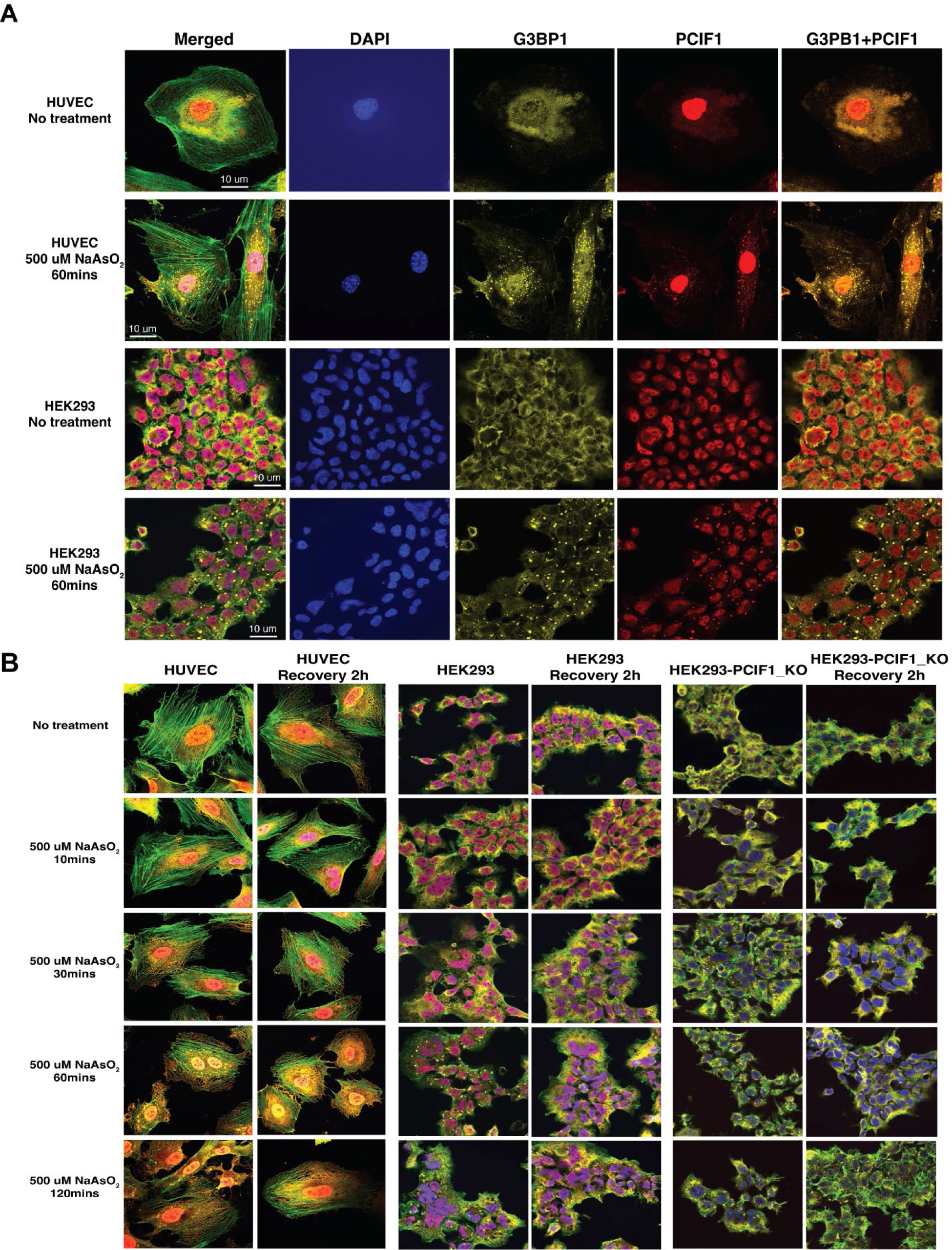
Indirect immunofluorescence shows co-localization of PCIF1 and G3BP1 in stress granules. HUVEC or HEK293 cells were grown on coverslips and treated with 500 µM NaAsO_2_ for the indicated times. Cells were fixed and stained with phalloidin (green), DAPI (blue), and antibodies targeting G3BP1 (yellow) and PCIF1 (red) as described in methods. (A) Representative images of each treatment showing the enrichment of PCIF1 and G3BP1 in stress granules upon NaAsO_2_ stress. (B) Cells were grown and treated with NaAsO_2_ for the indicated times. The NaAsO_2_ was removed, and cells were allowed to recover prior to fixing and staining. Images represent the consensus data from three independent biological replicates.

Notably, stress granules are temporary structures and disassemble when the stressor is removed^26^. We stressed HUVEC or HEK 293 cells for different times and assayed PCIF1 and G3BP1 localization after allowing them to recover for two (Fig 2B) or four hours (Fig S6). Indeed, stress granules dissipate over time as both the G3BP and PCIF1 foci are essentially absent from wild type cells two hours after the stressor is removed. We also performed these experiments in our PCIF1 knockout HEK293 cells and our data show that PCIF1 is not necessary for stress granule assembly or disassembly, but as shown in the PCIF1^-/-^ cells, disassembly is slowed in the absence of PCIF1 (Fig 2B, S6).

### PCIF1 physically associates G3BP and other stress granule component proteins

A search of several recent stress granule proteomics studies using different methods and baits showed that PCIF1 has not been detected as a stress granule protein prior to this work (Table S2, tabs 10-16)^27–31^. However, since PCIF1 behaves like a stress granule component by immunofluorescence, we wanted to learn if it directly interacted with other stress granule components (Fig 3). HEK 293 cells were subjected to standard (500 µM) or extreme stress (2.5 M NaAsO_2_) for 60 minutes, UV crosslinked, lysed, and PCIF was immunoprecipitated using PCIF1-targeting antibodies. Co-immunoprecipitated proteins were assayed by western blotting (Fig 3B). PCIF1 was able to pull down G3BP, a canonical stress granule component. We also used silver staining to examine if other proteins were immunoprecipitated by PCIF1 after one hour of standard stress. Multiple protein bands were recovered above background (Fig 3C, red stars).

**Fig 3.**
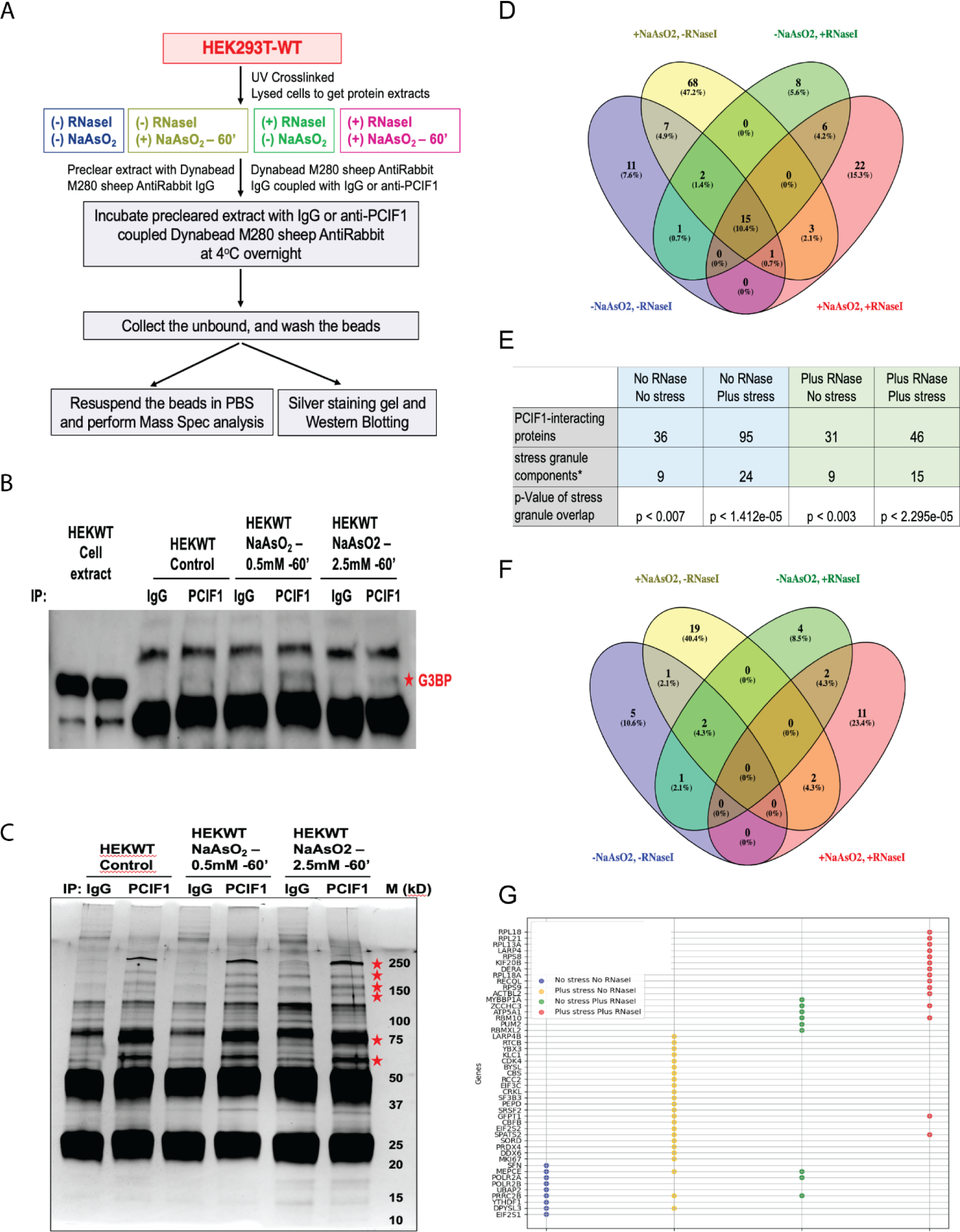
PCIF1 physically interacts with stress granule components. (A) Flowchart of PCIF1 protein interaction studies. (B and C) Cells were treated with the indicated dose of NaAsO_2_ for 60 minutes, UV crosslinked, lysed, and immunoprecipitated with a control (IgG) or PCIF1 antibody as indicated in methods. The resulting material was assayed by (B) Western blotting with a G3BP1-specific antibody or (C) SDS-PAGE and silver staining. (D-G) Immunoprecipitation experiments were repeated in the presence or absence of RNase I treatment. (D) A Venn diagram comparing the proteins co-purifying with PCIF1 (P-value < 0.05, log_2_FC > 25-fold enrichment over IgG control) in the presence or absence of RNase I. (E) The total number of proteins recovered in each IP and *the number of previously identified SG components are shown. (F) A Venn and (G) a line diagrams showing all SG components and the condition under which they were recovered. Representative images and data are shown from 3 independent biological replicates (panels B and C) or the average of three independent IP reactions (panels D-G).

Next, we used mass spectrometry to identify the PCIF1-interacting proteins in stressed and unstressed cells. Since PCIF1 is known to modify mRNAs and RNA is necessary core component of stress granules, we reasoned that performing the immunoprecipitation in the presence and absence of RNase could yield information about how PCIF1 interacts with other proteins before and during the stress response (Fig 3A) ^32^. Using a 25-fold enrichment over background and a P-value threshold of P <0.05, our data show that depending upon the condition, PCIF only interacts with a limited number of proteins, 36 or 31 ± RNase I respectively, in the absence of stress. Notably, the list of interacting proteins includes known PCIF1 binding partners such as POLR2A and POLR2B^12, 13, 33^. Further, as predicted by our localization studies, both nuclear and cytoplasmic proteins are immunoprecipitated in unstressed cells. After one hour of stress the repertoire of PCIF1-interacting proteins expands ∼2.5 fold (Fig 3D, Table S2).

Since PCIF1 redistributes to stress granules upon stress, we reasoned that we should examine if PCIF1 interacted with stress granule proteins. Next, we compared the lists of PCIF1-interacting proteins with lists of stress granule proteins obtained by different methods (Fig 3E, Table S2)^27–31^. Even in the absence of stress, PCIF1 physically associates with multiple proteins that are known to concentrate in stress granules (Fig 3E). As judged by a representation analysis, stress granule components are significantly (P-values from p < 0.041 to p < 1.589e-05 depending upon the comparison) enriched in nearly all samples and comparisons to previously published stress granule proteomics studies. When all proteins isolated from stress granules (a total of 1899) in several studies are taken into account, the PCIF1 interactome significantly (P-values from p < 0.007 to p < 1.412e-05) overlaps with stress granule proteins (Fig 3E) ^27–31^. Notably, the number of interactions with known stress granule proteins also increases ∼2 fold upon stress. Finally, fewer stress granule components (and proteins in general) interact with PCIF1 in the absence of RNA (Figs 3E-G), suggesting that some unknown RNA(s) are bridging these interactions.

### PCIF1 interacts with mRNA in normal and stress conditions

Since PCIF1’s protein interactome changes upon RNase treatment, our data suggested that PCIF1 bound directly to RNA. Indeed, this was consistent with prior reports showing that PCIF1 bound or modified mRNAs in the vicinity of the m7G cap^6, 33^. While cap-adjacent mRNA binding could conceivably be consistent with our data, we hypothesized that during stress PCIF1 might also interact with coding regions and 3’ UTR sequences like other writers of epitranscriptomic marks^1^. We used eCLIP with a PCIF1-targeting antibody to test this hypothesis in control and cells that have undergone either 10- or 60-minute exposure to NaAsO_2_ ^34^. eCLIP libraries from two independent biological replicates were compared and to minimize the inclusion of false positives, only shared mRNAs were retained. Our data show that PCIF1 directly binds to a small number (45) of mRNAs in the absence of stress, but that the number of mRNAs bound increases markedly to 120 and 767 mRNAs after 10 and 60 minutes of NaAsO_2_ respectively (Fig 4A, Table S3). A core group of 16 mRNAs were observed in all three conditions (Fig 4A). Further, with over 85% of signals coming from 3’ UTRs, coding sequences, non-coding exons, and distal introns, PCIF1 seemed to interact with multiple regions of the mRNAs in the absence of stress (Fig 4B). This interaction profile is very much consistent with PCIF1’s assumed co-transcriptional role.

**Fig 4.**
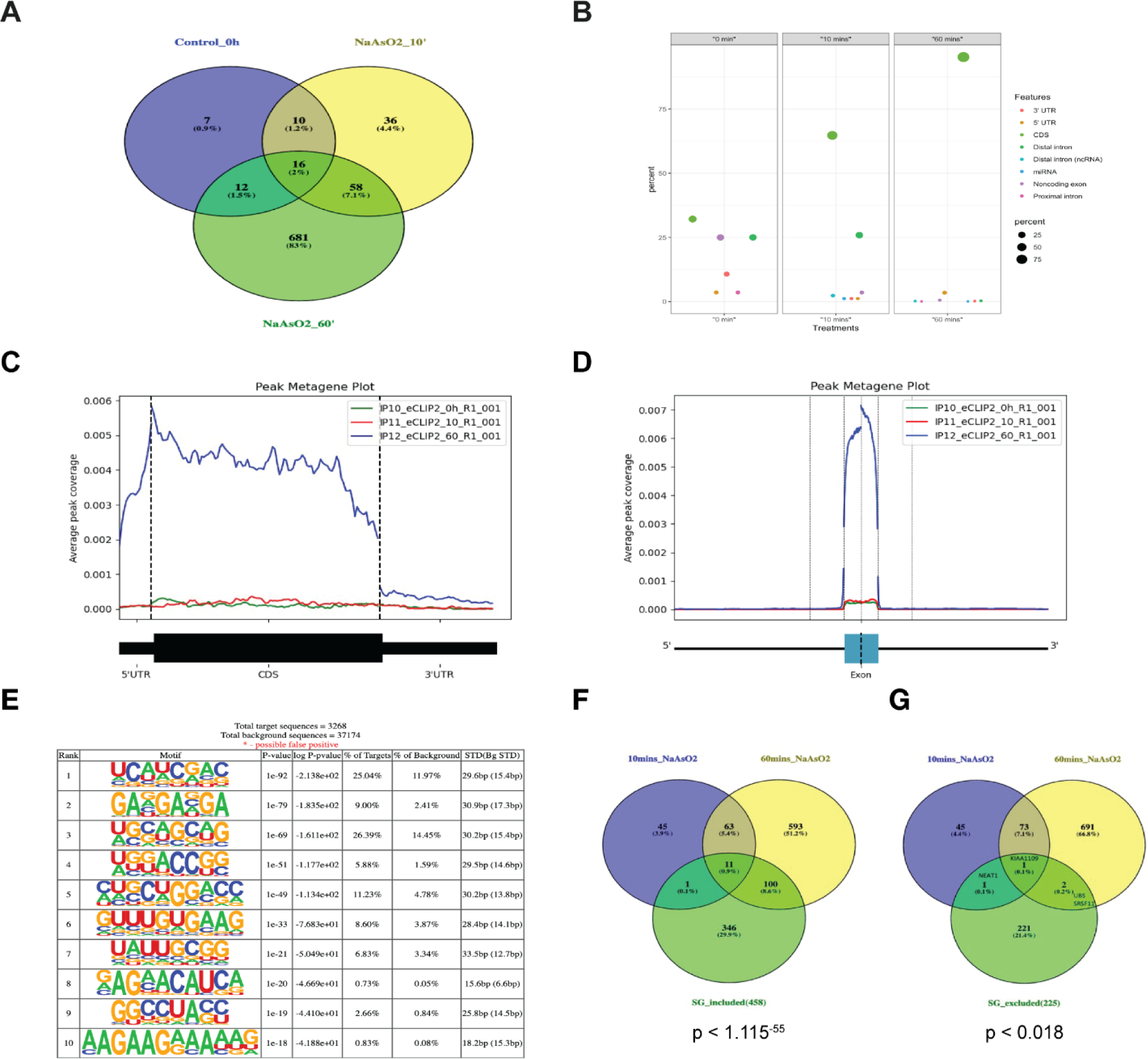
eCLIP experiments show that PCIF1 binds mature mRNAs are enriched in coding sequences during NaAsO_2_ stress. (A) A summary of the numbers of PCIF1-bound RNAs obtained via eCLIP (performed as described in methods) is shown for cells that were stressed with 500 µM NaAsO_2_ for 0, 10, or 60 minutes. (B) The mRNAs bound by PCIF1 were grouped by the feature of the mRNA where PCIF1 was crosslinked. The results are shown for 0, 10, and 60 minutes of stress. (C and D) PCIF1-bound sequences are enriched in the coding region of mature mRNAs during stress. (E) Sequence analysis shows possible consensus sequences for PCIF1-mRNA binding. (F and G) Our PCIF1-bound mRNAs recovered after 10 and 60 minutes of stress were compared to stress granule resident (F) or stress granule-excluded (G) mRNAs^23^. The eCLIP data shown are obtained from two independent biological replicates.

Much to our surprise, PCIF1’s mRNA binding profile both shifted and expanded markedly upon stress, with over 60% and 90% of binding corresponding to coding regions after 10 and 60 minutes of stress respectively (Fig 4C). Further, metagene plots show that PCIF1 bound almost exclusively within coding regions and exonic sequences of mature mRNAs after one hour of stress (Fig 4D). A sequence-based analysis of the crosslinked sequences recovered by eCLIP detected multiple possible PCIF1 recognition sequences (Fig 4E).

Next, since PCIF1’s protein interactome was significantly enriched in stress granule proteins, we compared our eCLIP data to mRNAs found in stress granules G3BP-labeled APEX-seq^23^. Ren and colleagues used nearly identical conditions (HEK293 cells, 500 µM NaAsO_2_, 60 minutes of stress) in their experiments. They identified core lists of 458 and 225 mRNAs that were significantly enriched in- or excluded from-stress granules respectively^23^. A direct comparison of our eCLIP-recovered mRNAs to these lists showed that after 60 minutes of stress, 111 of the 767 mRNAs in our PCIF1-eCLIP set were among the 458 mRNAs found in stress granules (Fig 4F, Table S3). Perhaps even more striking was only 3 of 767 were found among the stress granule-excluded mRNAs (Fig 4G).

Lastly, to examine if our observations can be explained by a co-transcriptional process, we performed ChIP-Seq using antibodies targeting PCIF1, and chromatin modifications that mark transcription start sites (Histone H3 with acetylation on lysine 9, (H3K9)), gene bodies (Histone H3 with trimethylation on lysine 36, (H3K36me3)) in stressed and unstressed cells. As predicted and consistent with its known role in methylating cap-adjacent adenosines, PCIF1 showed an enrichment near the transcription start site in unstressed cells but, consistent with transcriptional shutdown of the depicted genes, ChIP-seq signal was lost in stressed cells (Figs 5, S11). The decrease in PCIF1 ChIP-Seq occurs simultaneously with an increase in PCIF1’s binding to its mRNA targets (Figs 5, S11).

**Fig 5.**
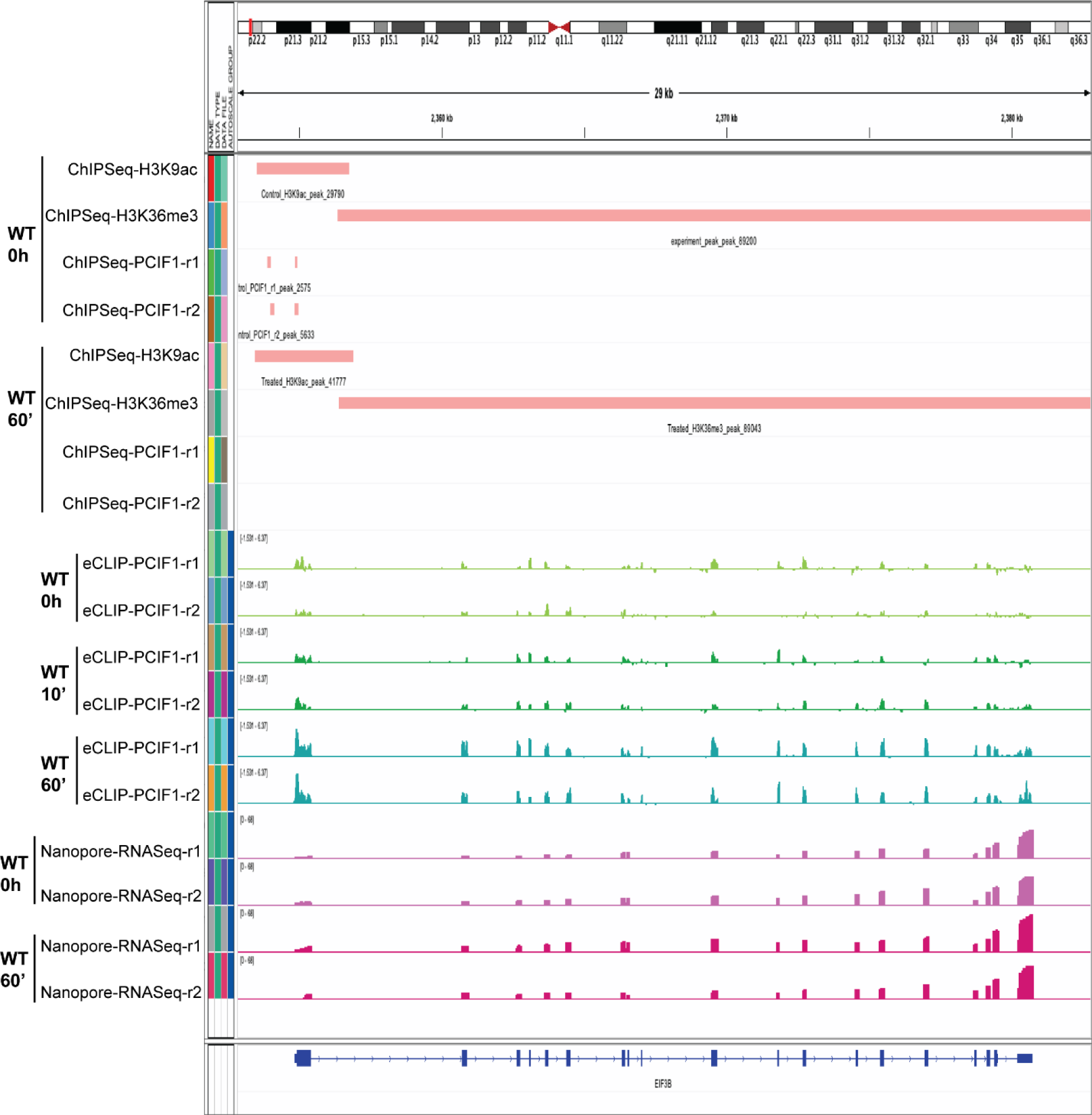
Multiple sequencing techniques show PCIF1’s association with mRNA is not co-transcriptional. Visual representation of ChIP-Seq (IP antibodies were for Histone H3K9ac, Histone H3K36me3, and PCIF1), eCLIP (with PCIF1 antibody), and Oxford Nanopore direct RNA sequencing profiles mapped to EIF3B mRNA, a representative PCIF1-bound, stress granule-associated mRNA. Data are shown from both non-stressed and stressed WT cells. The data shown are the consensus of ChIPSeq, eCLIP and direct RNA sequencing results from duplicate experiments. The plot was made with IGV.

### Oxford Nanopore long read direct mRNA sequencing reveals negligible effects of PCIF1 knockdown on global mRNA levels in both stressed and unstressed cells

We used oxford nanopore direct RNA sequencing to assess the changes in bulk mRNA levels when comparing WT and PCIF1^-/-^ HEK293 cells. Prior work had shown that a small population of mRNAs were changed substantially when PCIF1 was deleted. Our data concur with those earlier observations, and we see only a handful of mRNAs were changed in our comparisons (Fig 6, Table S4). Finally, we did observe changes in the levels of some mRNAs when comparing unstressed and stressed cells (Fig 6, Table S4).

**Fig 6.**
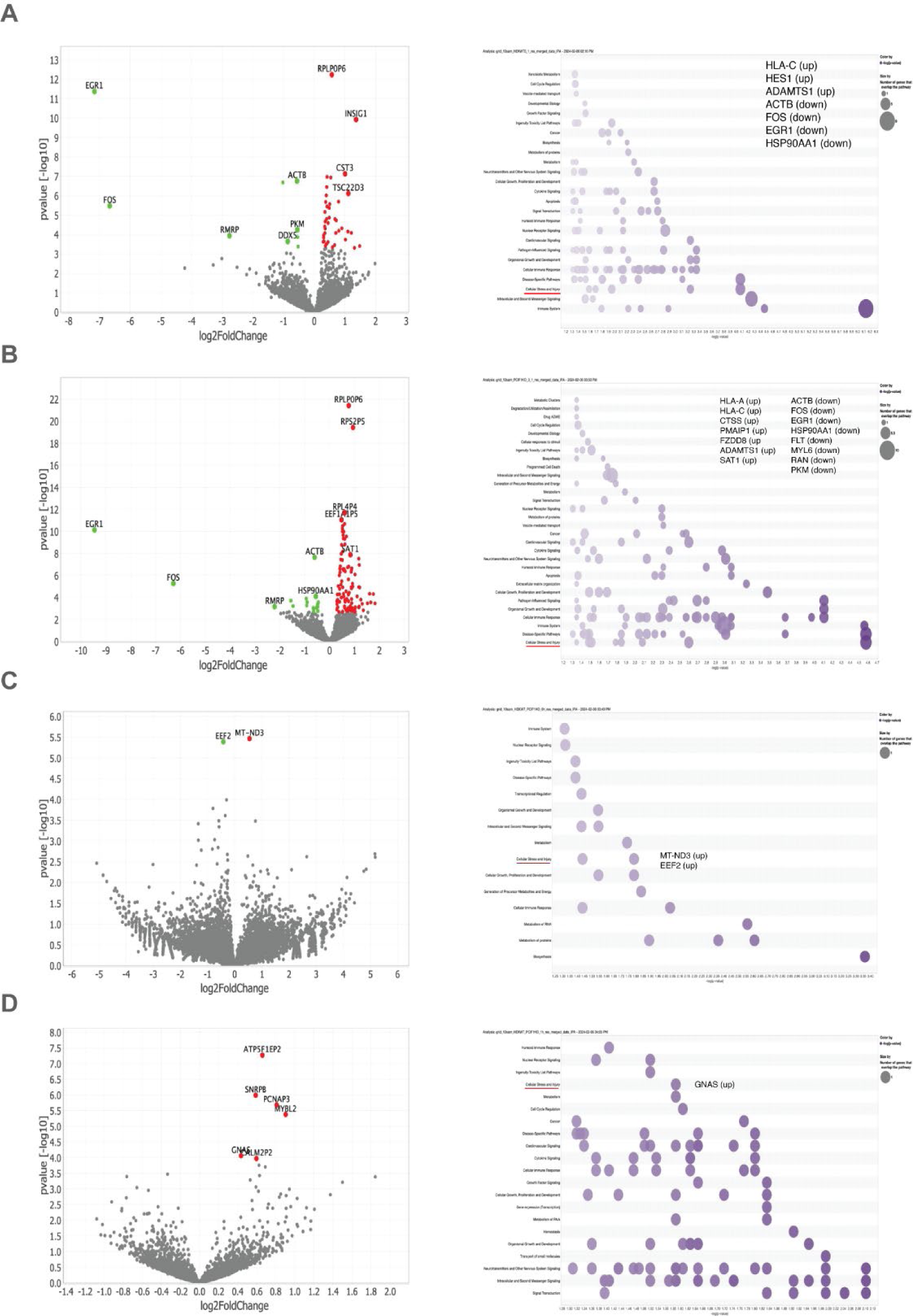
Oxford Nanopore direct RNA sequencing of non-stressed and stressed WT and PCIF1^-/-^ HEK293 cells. WT or PCIF1^-/-^ HEK293 cells were stressed with 500 µM NaAsO_2_ or left unstressed for 60 minutes. Total RNA was harvested from cells and Oxford Nanopore direct RNA sequencing was performed as indicated in the methods. Changes in transcript abundance were mapped by pairwise comparisons. Left column: Changed RNAs are shown for (A) unstressed WT cells vs stressed WT cells, (B) unstressed PCIF1^-/-^ cells vs stressed PCIF1^-/-^ cells, (C) unstressed WT cells vs unstressed PCIF1^-/-^ cells, (D) stressed WT cells vs stressed PCIF1^-/-^ cells. Right column: GO term analysis of the increased/decreased genes is shown. The data shown are the consensus of direct RNA sequencing results from duplicate experiments.

## Discussion

Perhaps the four most salient findings of our study are that: (1) a population of PCIF1 is always present in the cytoplasm, (2) PCIF1 actively re-distributes to stress granules and interacts with stress granule proteins upon stress, (3) PCIF1 binds directly to mRNA coding regions, and (4) the mRNA binding profile of PCIF1 greatly expands and includes stress granule resident – but not stress granule excluded – mRNAs during the stress response.

Our work is consistent with all other prior PCIF1 studies and shows that PCIF1 is predominantly nuclear; however, as shown elsewhere, we show that at least a population of PCIF1 is cytoplasmic (Fig 1)^24, 25^. PCIF1’s sustained presence in the cytoplasm opens up new possible roles in post-transcriptional gene regulation. We present three distinct lines of evidence that PCIF1 has a role in stress granules. First, our immunofluorescence data show PCIF1 localizing to stress granules upon stress onset and dissipating from them upon recovery (Figs 2, S4-S7). Second, our proteomics data also show that PCIF1 interacts with a significant number of stress granule proteins (Fig 3, S8). Finally, our eCLIP data show that PCIF1-bound mRNAs are significantly enriched in mRNAs that aggregate in stress granules (Fig 4). Collectively, these data are consistent with PCIF1 having a functional role in stress granules. Moreso, the observations that stress granule formation is hindered by the overexpression of PCIF1, and that stress granule dissolution is delayed in PCIF1 knockout (Fig 2B, S6, S7) cells offers further evidence that PCIF1 has a functional role within stress granules. In fact, those data are consistent with PCIF1 having an active role in stress granule disassembly.

Notably, our IP-Mass spec data, even when only considering unstressed cells, are markedly different than a previously published study which used the same parent HEK293 Trex-Flip in cell line^33^. Importantly, by using UV crosslinking cells prior to lysis and immunoprecipitating PCIF1, our methodology was substantially different than the BioID fusion protein strategy employed the earlier work ^33^. Furthermore, since we focused on the cytoplasmic function(s) of PCIF1, we fractionated our cell lysates and performed our IP’s using nuclear-depleted cytoplasmic extracts as opposed to the whole cell lysates used previously^33^. Our fractionation isn’t perfect as ∼10% of our interacting proteins are nuclear including both POLR2A and POLR2B in unstressed cells. Further, as nearly 100% of cytoplasmic PCIF1 aggregates in stress granules, we wouldn’t expect much overlap between proteins recovered from our stressed cells and those recovered via BioID-labeling and pulldown obtained from unstressed cells^33^.

Perhaps our most surprising findings were that PCIF1 physically interacted with a large number of mRNAs, and that during stress, these interactions were almost exclusively with the coding regions of the mRNAs (Fig 4). This is a stark contrast with expectations of 5’ UTR-focused binding, based on the enrichment of PCIF-deposited m^6^Am as assayed by m6Am-Exo-Seq^7^. Our data show a core list of 16 transcripts that were ever-present in stressed and unstressed cells, but that profile essentially triples upon NaAsO_2_ stress. Perhaps more strikingly, when we compared our PCIF1-bound mRNAs to those obtained using G3BP-APEX-Seq, a significant percentage of the 767 mRNAs in our eCLIP data had been demonstrated to aggregate in stress granules during NaAsO_2_ stress^23^. Conversely, only 3 mRNAs detected through eCLIP were categorized as ‘stress granule-excluded’ by the same measure^23^.

We are currently working to determine both the mechanism and consequence of PCIF1:mRNA association during stress. Regarding the first question, our initial hypothesis is that the previously identified RNA binding region that PCIF1 uses to coordinate the formation of cap-adjacent m^6^Am is responsible for mRNA binding^6^. Another possibility is that a strongly positively charged face of the protein adjacent to PCIF’s methylation active site enriched in arginine residues (Fig S13) serves as the interface for RNA binding^6^. Still a third possibility is that a novel portion of the protein (or interacting partner) is crucial for PCIF’s ability to bind mRNAs.

Regardless of the physical parameters of how PCIF1 interacts with mRNAs during stress, the functional consequence of that interaction is of keen interest. Collectively our observations strongly suggest that PCIF1 has functions beyond co-transcriptional cap-adjacent methylation of adenosines. Intriguingly, in vitro studies have shown that PCIF1 can methylate RNA oligonucleotides both internal adenosines and 5’ end adenosines, even in the absence of an m^7^G cap^35–37^. With this information in mind, the interactions we show raise the possibility that PCIF1 binds and modifies mRNAs by adding m^6^Am internally - within their coding regions - during stress. Further, multiple studies have shown that mRNAs dynamically enter and exit stress granules during the stress response, but the gatekeeping mechanism remains unknown^21, 22, 38^. A provocative hypothesis is that PCIF1, either by the addition of m^6^Am or an as of yet unidentified function, could contribute to this gatekeeping process, likely by facilitating the exit of selected mRNAs from stress granules.

## Supporting information

Supplement Table 1. Key resources table

Supplement Table 2. PCIF1 Mass Spectrometry results and Comparisons to known protein components of stress granules

Supplement Table 3. PCIF1 eCLIP results and Comparisons to mRNAs known to localize to or be excluded from stress granules

Supplement Table 4. Changed mRNAs observed in WT and PCIF1-/- cells as observed by Oxford Nanopore direct RNA sequencing

## Author contributions

TTT and DLK conceived the studies and designed the experiments. TTT performed the bulk of the cell culture, cell fractionation, immunoprecipitation, immunofluorescence, RNA extraction experiments, prepared the libraries for Oxford Nanopore Direct RNA Sequencing, and performed the bioinformatics analysis and generated the resulting plots. SR aided with developing the data analysis pipeline for Oxford Nanopore Direct RNA Sequencing. TLW assisted with cell culture and western blotting and mined the Human Genome Atlas to determine the localization of PCIF1-interacting proteins. SK performed literature searches and assisted with the bioinformatics analysis to calculate the overlap of stress granule components and PCIF1-interacting proteins. SYJ performed and analyzed the data generated by the mass spectrometry studies. All authors have read and approved the manuscript.

## Acknowledgements

We would like to thank the staff of the Advanced Cellular and Tissue Microscopy Core Facility at the Houston Methodist Research Institute(HMRI) for technical support. We thank Dr. Abhinav Jain and the MDACC Epigenomics Profiling Core Facility for their assistance with the ChIP-Seq assays. This work was supported by start-up funds provided by the HMRI and a grant from the National Institutes of Health (1R35GM137819 both to DLK). The content presented here is solely the responsibility of the authors and does not represent the official views of the HMRI, Baylor College of Medicine, or the National Institutes of Health.

## Methods

### Key Resources

All vendors and catalog numbers for antibodies, enzymes, reagents, kits, other assorted consumables, and software versions are recorded in Table S1.

### Cell culture

HUVEC, U2OS, and HEK293 Flp-In T-REx cells were maintained under 5% CO2 at 37°C. HUVEC cells were cultured in EBM^TM^ Basal Medium supplemented with 10% FBS, 1% penicillin/streptomycin, and EGM^TM^ Endothelial Cell Growth Medium SingleQuots Supplements. U2OS was cultured in McCoy’s 5A medium supplemented with 10% FBS. HEK293 Flp-In T-REx was cultured in DMEM high glucose medium supplemented with 10% FBS, 1% penicillin/streptomycin, L-glutamine, 100 ug/ml Zeocin, 15 ug/ml Blasticidin. For cell passaging, cells were first washed with PBS. 0.25% Trypsin-EDTA was added to the dishes, which were then incubated at 37°C for 2-5 min. FBS containing medium was added to inactivate trypsin and the cells were collected by centrifuging at 2,000 rpm for 3 min. Cells were counted and split as needed for different purposes. All cell cultures are routinely monitored (and tested negative for) mycoplasma contamination using the MycoStrip^TM^ – Mycoplasma Detection Kit.

### Cell transfection

For transfection, a mammalian expression plasmid encoding the human tagged PCIF1 (NM_022104) ORF clone) were amplified in E. coli GC5 DH5α and extracted using Zymo Maxiprep kits. Purified plasmids were transfected into HUVEC and HEK293 cells using the Lipofectamine 3000 Transfection reagent according to the manufacturer’s instructions. After transfection, the cells were either lysed for Western Blotting or fixed to perform immunofluorescence procedure.

### Cell fractionation

Cell fractionation was performed with slight modifications to an earlier protocol^39^. Cultured cells were washed in ice-cold PBS pH 7.4, placed on ice, and scraped from the dish using a cell scraper and collected in 1.5 ml micro-centrifuge tubes in 1 mL of ice-cold PBS. Cells were pelleted for ∼10 sec in an Eppendorf table top microfuge and the supernatant was removed from each sample. Cell pellets were resuspended in 900 μl of ice-cold 0.05% NP40 in PBS supplemented with Protease Inhibitor Cocktail, Phosphatase Inhibitor II, Phosphatase Inhibitor III, and 0.05 mM PMSF. 300 μl of the lysate was taken as a whole cell extract. The remaining lysate was centrifuged for 10 sec at maximum speed and the supernatant was collected as the cytoplasmic fraction. The cell pellets were resuspended in 140 μl lysis buffer and designated as “nuclear fraction”. The nuclear fractions were sonicated in SDS containing lysis buffer with the settings as “time on/off” to 30sec, cycle number: 60, run on “low” for 1 hour at 4°C. After sonication, the samples were centrifuged for 15 mins at 4°C at max speed and supernatant was collected for western blotting.

### CRISPR knockout cell line

Two clonal PCIF knockout HEK293 Flip-in Trex cell lines (clones H5 and G5) were made by Synthego using their standard CRISPR-Cas gene knockout pipeline^40^. The guide RNAs used are presented in Fig S2.

### Western blotting

Western blots were performed essentially as described earlier^41^. Cell lysates were prepared as above, or cell pellets were washed with ice cold PBS and incubated on ice in M-PER lysis buffer supplemented with Protease Inhibitor Cocktail, Phosphatase Inhibitor II, Phosphatase Inhibitor III, and 0.05 mM PMSF. Proteins were separated by electrophoresis using pre-cast SDS-PAGE gel and transferred to a PVDF membrane. The membrane was blocked in 5% nonfat milk and incubated with primary antibody overnight at 4°C. The membranes were washed three times with 1x TBST and incubated with HRP-conjugated antibody for 2 hours at room temperature. The protein bands were detected by western ECL Substrate, visualized using ChemiDoc MP Imaging System, and quantified by Biorad ImageLab software. We performed three independent replicates and representative blots are shown. Nine antibodies were used to detect the PCIF1 expression levels (Supplement Table 1), targeting the entire protein as well as N-terminal and C-terminal region of PCIF1 protein, respectively.

### Immunofluorescence staining

HUVEC and HEK293T cells were grown on fibronectin coated coverslips. Cells then were treated with 500 μM NaAsO_2_ for 0, 10, 30, 60, or 120 mins. Cells then were fixed with 4% PFA for 20 mins at 4°C. PFA was removed and cells were washed twice with PBS and permeabilized with blocking/permeabilization buffer (0.1% Triton X-100 and 2% BSA in PBS) for 1 hour at 4°C. Primary antibody solution was prepared in blocking/permeabilization buffer (1:100) and added to the wells after removing the initial blocking buffer. The samples were incubated at 4°C overnight, followed by incubation with Alexa Fluor 555 or Alexa Fluor 647 conjugated goat anti-mouse or anti-rabbit secondary antibodies for 1 hour at room temperature. After hybridization with secondary antibodies, the wells were washed twice with PBS and incubated with F-actin Alexa Fluor 488 phallodin for 30 mins at room temperature. Coverslips were mounted onto slides using Diamond antifade Mountant with DAPI.

An Olympus FV3000 confocal microscope was used to capture images of samples using the FV31S-SW (Ver.2.6) acquisition software. The system utilized a combination of Cooled GaAsP and Multi-Alkali photomultiplier detectors and a Galvano scanner to acquire images. The 100XS UPLSAPO objective lens was used to acquire images at a 0.124 um/pixel (1024×1024) resolution and pinhole of 1.0 Airy Disk (388um confocal aperture). Diode lasers (405nm, 488nm, 561nm, 640nm) were used to excite the samples and capture the emission signal from DAPI, F-actin, G3BP1, and PCIF1, respectively.

### Sucrose gradient centrifugation for polysome analysis

Cells were fractionated using linear 10-50% sucrose gradients as described earlier ^42^. Briefly, cells underwent treatment with 100 ug*/*ml Emetine dihydrochloride for 15 min at 37°C, washed twice with PBS containing Emetine (100 ug*/*ml) and 0.01M MgCl_2_. Cells were then scraped off the dish and pelleted by centrifugation at 2000 rpm for 3 minutes. All further steps were carried out on ice using pre-chilled centrifuges. Cell pellets were lysed using five volumes of ice cold lysis buffer (50 mM Tris-HCl, pH 7.5, 10 mM KCl, 10 mM MgCl_2_, 150 mM NaCl, 1% Triton X100, 2 mM DTT, 0.5 mM PMSF, 100 ug*/*ml Emetine, Protease Inhibitor Cocktail, Phosphatase Inhibitor II, Phosphatase Inhibitor III, and RNase-In Plus). The lysates were incubated on ice for 30 min with gentle mixing every 10 minutes. Centrifugation at 20,000 rpm for 15 min removed nuclei and other debris. The clear supernatants were then applied to 10–50% linear sucrose gradients. These gradients were centrifuged for 3 h at 35,000 rpm in a SW41-Ti rotor at 4°C. Fractions of 0.9 ml were collected from the bottom and the absorbance at 254 nm was continuously monitored using Brandel BR 188 Gradient fractionation system.

### Immunoprecipitation

Cells were UV crosslinked at 254-nm using a UVP CL-1000 Ultraviolet Crosslinker with the energy setting of 400 mJoules/cm^2^ and extracted the cytoplasmic fraction using ice-cold 0.05% NP40 in PBS supplemented with Protease Inhibitor Cocktail, Phosphatase Inhibitor II, Phosphatase Inhibitor III, and PMSF 0.05 mM. Antibodies were used in these experiments as PCIF1 (N-terminal) Rabbit and IgG-Rabbit. Dynabead (M280 Sheep anti-Rabbit) was washed twice with lysis buffer and resuspended in 100 μl buffer. The washed beads were pre-bound to 10 μg of the antibody by continuous rotation for 2 hours at room temperature. Then, antibody-coupled beads were washed twice and incubated with cytoplasmic extracts overnight at 4°C. Beads were then washed three times with high salt buffer and twice with no salt buffer; and eluted in 1X Western buffer at 65°C for 10 mins. High salt, no salt and 1X Western Buffer were provided in the IP Antibody Validation Kit from EclipseBio. The bound and unbound samples were loaded onto 4-15% Biorad gel and analyzed using either western blotting, silver staining, and/or mass spectrometry.

### Silver staining

After IP, the unbound and bound samples was loaded onto 4-15% Biorad gel; and stained using Pierce Silver Stain for Mass Spectrometry Kit. The gel was washed twice for 5 min in ultrapure water. Next, water was removed; and the gel was incubated in fixing solution (30% ethanol, 10% acetic acid) for 15 min at room temperature. The fixing step was repeated after replacing solution. Then the gel was washed with 10% ethanol twice for 5 min each following up with washing in water twice for another 5 min each. In the last step, the gel was briefly incubated in Silver Stain Sensitizer and developed with Silver Stain Developer in 2-3 min until protein bands appeared. When the desired band intensity is reached, we replaced developer working solution with Stop solution (5% acetic acid). The gel was washed briefly, then replaced with acetic acid or water for storage.

### Mass Spectrometry

PCIF1 protein complex identification was processed as described previously with the following modifications^43^. First, the purified PCIF1 complex underwent digestion on beads using 1 μg of Trypsin/Lys-C enzyme overnight, then the digested peptides were subjected to enrichment and desalting utilizing an in-house STAGE tip column packed with 2 mg of C18 beads (3 µm, Dr. Maisch GmbH, Germany) and vacuum dried^44^. The resuspended peptides were subjected to analysis using Orbitrap Fusion mass spectrometers coupled with an Easy-nLC 1000 nanoflow LC system. Nano-HPLC separation was carried out using an in-house trap column (2 cm × 100 µm i.d.) and a 5 cm × 150 µm capillary separation column packed with 1.9 µm Reprosil-Pur Basic C18 beads (Dr. Maisch, r119.b9.) employing a discontinuous gradient of 2–26% acetonitrile, 0.1% formic acid at a flow rate of 800 nl/min. The Mass spectrometry analysis was conducted in a data-dependent mode, acquiring fragmentation spectra of the 30 strongest ions under the control of Xcalibur software (Ver.4.1). The parental ion was acquired in the Orbitrap with a full MS range of 300–1400 m/z at a resolution of 120,000. Fragmented MS/MS spectra were acquired in the ion-trap using higher-energy collisional dissociation (HCD) in rapid scan mode. Subsequently, the MS/MS spectra were searched against the target-decoy Human RefSeq database (release Jan. 21, 2020, containing 80,872 entries) utilizing the Mascot algorithm (Ver.2.4), within the Proteome Discoverer (Ver.2.1) interface. The search parameters allowed for a precursor mass tolerance of 20 ppm and a fragment mass tolerance of 0.5 Da, with up to two maximum missed cleavages permitted. Dynamic modifications including acetylation of N-term and oxidation of methionine were considered. The assigned peptides were filtered with a 1% false discovery rate (FDR) using Percolator validation based on q-value. Peptide Spectrum Matches (PSMs) output from PD (Ver.2.1) was utilized to group peptides onto the gene level employing the ’gpGrouper’ algorithm (Ver. 1.0)^45^. An in-house program, gpGrouper, applies a universal peptide grouping logic to accurately allocate and provide MS1-based quantification across multiple gene products. Gene-protein products (GPs) quantification was performed using the label-free, intensity-based absolute quantification (iBAQ) approach. Missing values in the mass spectrometry recovery were replaced with half of the minimum iBAQ of the entire dataset. After log2 transformation of the dataset, differential analysis (t test) was performed.

### Stress granule component comparisons

Several manuscripts were used as starting point to determine the stress granule proteome^27–31^. The overlap of stress granule proteins and PCIF1-interacting proteins was determined for the individual studies and a collected aggregate list of 1899 stress granule proteins. The representation factor and significance of the overlap was determined using the following webtool: <u>Statistical significance of overlap of two groups of genes</u> (nemates.org). The precise mathematical formulas used for these calculations are listed here: http://nemates.org/MA/progs/representation.stats.html. The values for proteins in each list are shown in Table S2. Further, the total number of genes in the human genome was set to 19,062 per this paper ^46^.

### eCLIP

Briefly, HUVEC cells were cross-linked on ice with UV irradiation (254 nm) at 400 mJ cm^−2^ using a UVP CL-1000 Ultraviolet Crosslinker. Cells were then collected by centrifugation, lysed and cell lysates were frozen at −80°C until further processing by EclipseBio. The subsequent immunoprecipitation (PCIF1 antibody), library preparation, sequencing, and quality control testing were performed by using the standard eCLIP pipeline from EclipseBio.

### Analysis of eCLIP data

Raw sequencing reads were demultiplexed by removing UMIs from the sequencing reads and appended to the read name for PCR deduplication. Both 5’ and 3’ adapters were trimmed off using Cutadapt software (Ver.3.2). These trimmed reads were subjected to FastQC (Ver.0.11.9) for quality control. The quality threshold was set to 30. Trimmed reads were mapped to a repetitive element database using STAR rmRep to remove repetitive elements and help prevent any spurious artifacts from rRNA and other repetitive reads. A run of FastQC using output from STAR rmRep was performed to verify the read quality before doing STAR genome mapping with human genome GRCh38 (hg38). Bam files were sorted and indexed using SAMtools (Ver.1.9). The output from the SAMtools view generated bigwig files, which were used to visualize the data using IGV software (Ver.2.17.4). eCLIP peaks were called using CLIPper (Ver.1.3.0) (EclipseBio, proprietary). All Venn Diagrams were made with the Venny webtool (Ver.2.0)^47^.

### Chromatin immunoprecipitation sequencing (ChIP-Seq)

HUVEC cells underwent cross-linking with 1% formaldehyde adding directly to the media and gently shaking at room temperature for 8 minutes for histone modifications or 15 minutes for PCIF1. Then, glycine at a final concentration of 0.125 M was added to the media with slow shaking for 5 minutes at room temperature to stop the cross-linking. Cells were then washed twice with ice-cold PBS, pelleted, and stored at −80°C. The subsequent steps of the chromatin immunoprecipitation (ChIP) assays were performed at the Epigenomics Profiling Core, MD Anderson Cancer Center. Nuclei were isolated from cross-linked cells followed by chromatin preparation and sonication to obtain fragment size ranging from 200-600bp. Chromatin lysate equivalent of 5 million cells were subjected to chromatin immunoprecipitation with antibodies specific to PCIF1, H3K9ac, and H3K36me3 antibodies. ChIPs were performed as described previously with some modifications^17, 48^. Input and ChIP DNA libraries were prepared using NEB Ultra II DNA library prep kit following manufacturer’s protocol and subjected to Next-generation sequencing on an Illumina NextSeq 500 to obtain 20-30 million 75bp single reads per sample.

### Analysis of ChIP-Seq data

FASTQ files were subjected to FastQC for Phred quality score of 30 or more. Reads were mapped to a repetitive element database to remove repetitive elements and help prevent any spurious artifacts from rRNA and other repetitive reads. The output was verified using FastQC for read quality before the BWA genome mapping with human genome hg38. Bam files were sorted and indexed using SAMtools. The peak calling was performed using MACS2 (ver.2.1.2) and converted to BED files which were used to visualize on IGV.

### Nanopore direct RNA Sequencing

Total RNA was harvested from cells using Direct-Zol RNA purification kits, and the integrity of purified RNA was assayed using a TapeStation 4200. The RNA library preparation was performed using 2ug total RNA extracted from HEK293 wildtype and PCIF1-KO cells using Nanopore Direct RNA Sequencing kit SQK-RNA002 according to the manufacturer’s instructions. The libraries were loaded to GridION flow cells (R9.4.1) and sequenced using Nanopore GridION X5.

### Analysis of Nanopore RNA Directed Sequencing data

Data acquisition and real-time basecalling were carried out during sequencing run by MinKNOW software (Ver.1.4.2). The fast5 and fastq output files were retrieved and subjected to FastQC and NanopPot (Ver.1.42.0) for quality control. The fastq reads were mapped with human transcriptomics using Minimap2 (Ver.2.28) and htseq-count command from the HTSeq (Ver.0.12.4) was used to preprocess data for differential expression analysis by counting the reads. Differential expression analysis was carried out using DESeq2 (Ver.1.43.5). The plots were generated in the R suite (Ver.4.4.0) using Bioconductor software (Ver.3.19). Unique mapped reads were visualized on IGV (Ver.2.17.4).

### Statistics and reproducibility

At least three biological replicates were utilized for each experiment unless specified otherwise. The results were reported as the mean ± standard error of the mean (SEM) or standard deviation (SD) as indicated in each legend and table. Statistical significance between groups were evaluated using t-test, and *P* value was indicated in the figure legends and tables. Pearson correlation coefficient (PCC) was calculated to determine correlations. All the western blotting, IP, IF, Mass Spec, eCLIP, ChIP-Seq, and Nanopore Direct RNA sequencing experiments used two (eCLIP, some Nanopore Direct RNA Seq) or three independent biological replicates with consistent results.

### Data availability

The sequencing data supporting the findings of this study have been submitted to the Gene Expression Omnibus (GEO) database and the accession code is pending. The original and processed data are provided as supplementary materials. All additional data that support the findings of this research can be obtained from the corresponding author on reasonable request.

---

**Supplementary Fig S1.**
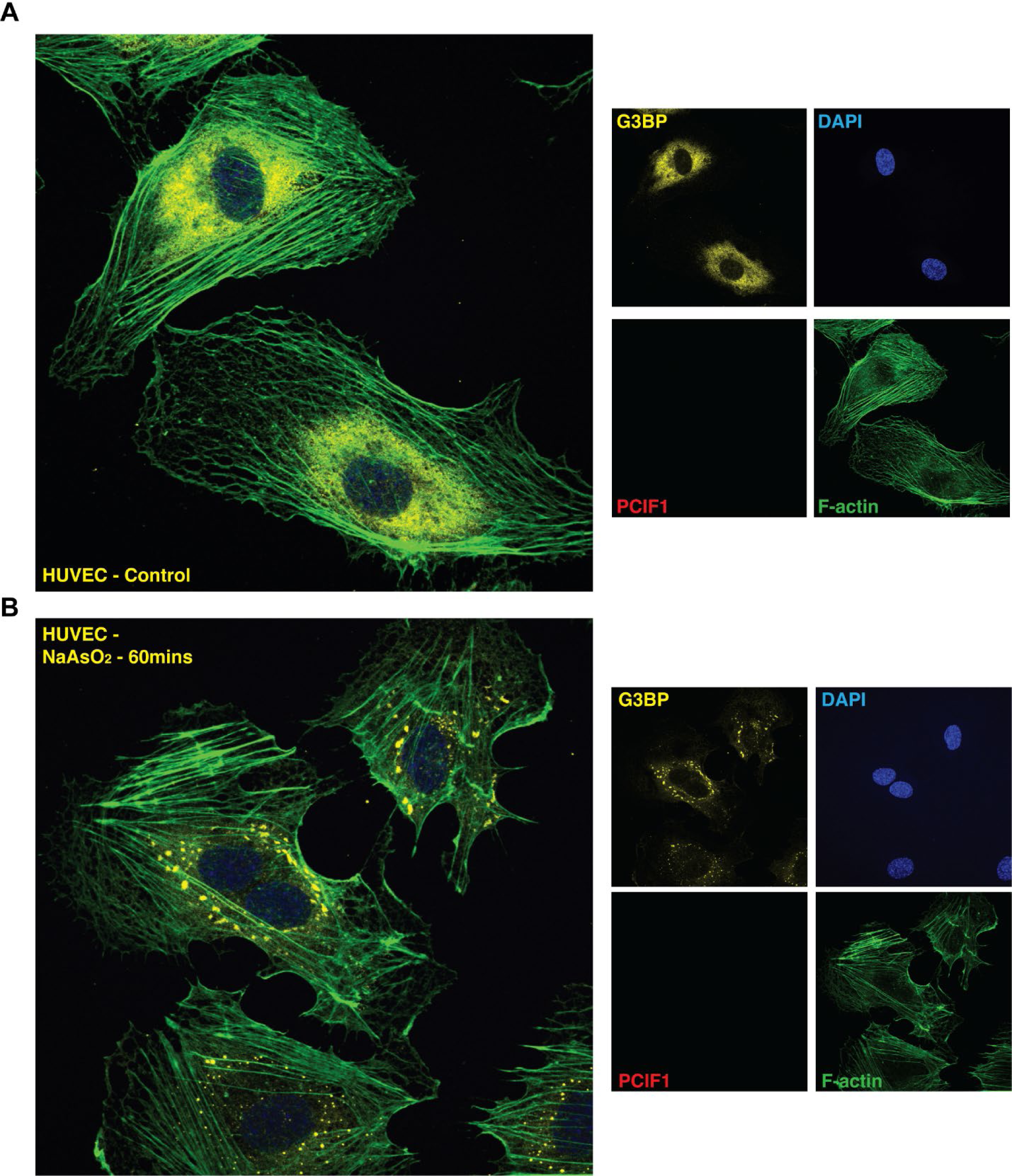
Control experiments to assess the background signal of mouse and rabbit secondary antibodies show minimal background signal. HUVEC cells were grown on coverslips and either (A) mock treated or (B) stressed with 500 µM NaAsO_2_ for 60 minutes. Cells were fixed and stained with G3BP-targeting antibodies in addition to DAPI and phalloidin. The stained cells were then probed with either mouse (A) or rabbit (B) secondary antibodies, without using primary PCIF1-targeting antibodies. Representative images are shown from three independent biological replicates.

**Supplementary Fig S2.**
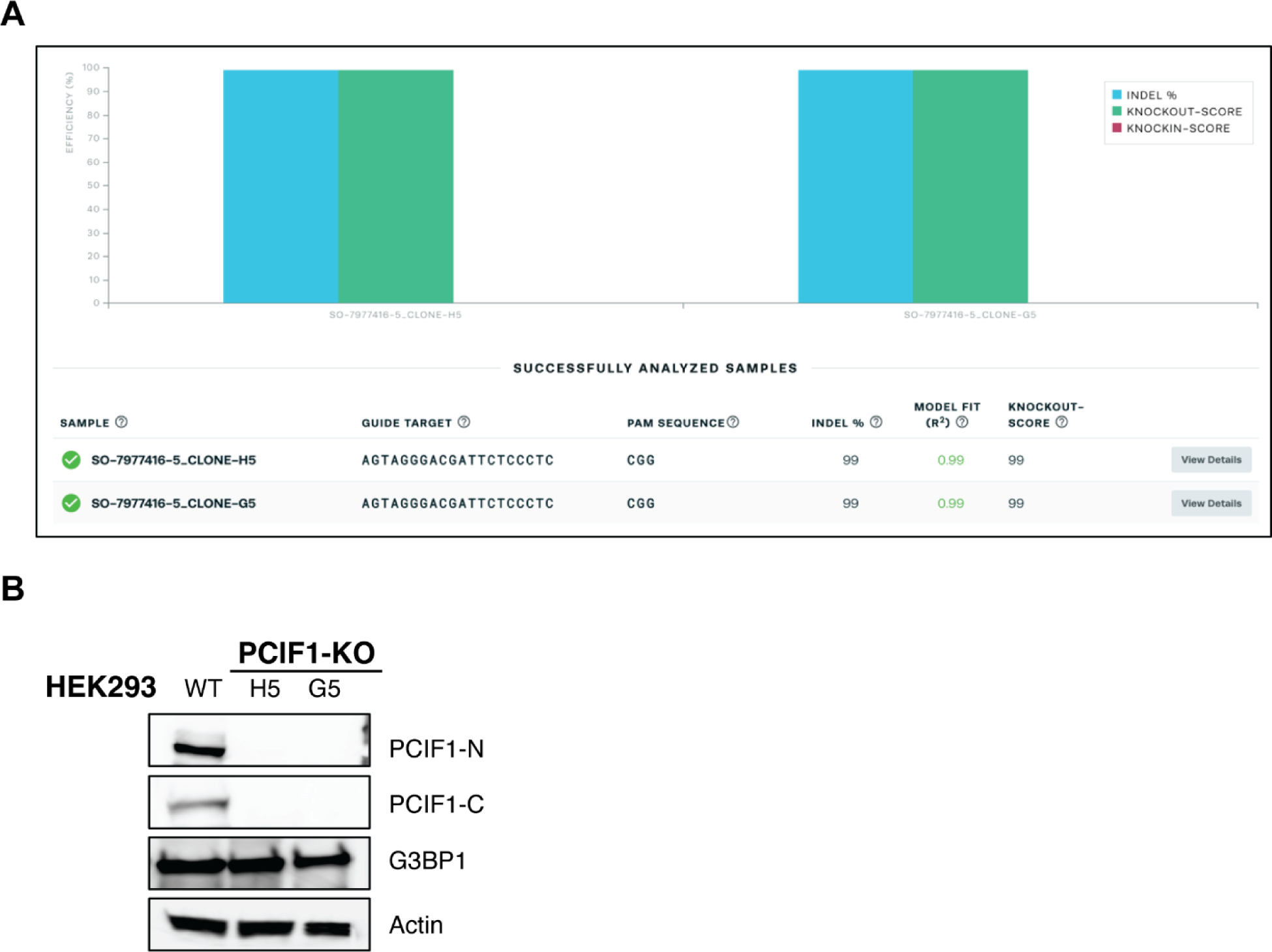
PCIF1 was effectively deleted from HEK293 cells using CRISPR-Cas technology. (A). Estimation of PCIF1 knockout efficiency for two independent clonal PCIF1^-/-^ HEK293 knockout cell lines. The guide RNA target sequences are also indicated. (B) Western blots using cell lysates harvested from WT and PCIF1^-/-^ HEK293 cells (H5 and G5 clones) were used to assess the efficiency of the knockout. Two distinct antibodies targeting PCIF1 (one targeting the N-terminus and one targeting the C-terminus) were used. The blots were also probed for Actin and G3BP1 as loading controls. Representative images are shown from 3 biological replicates.

**Supplementary Fig S3.**
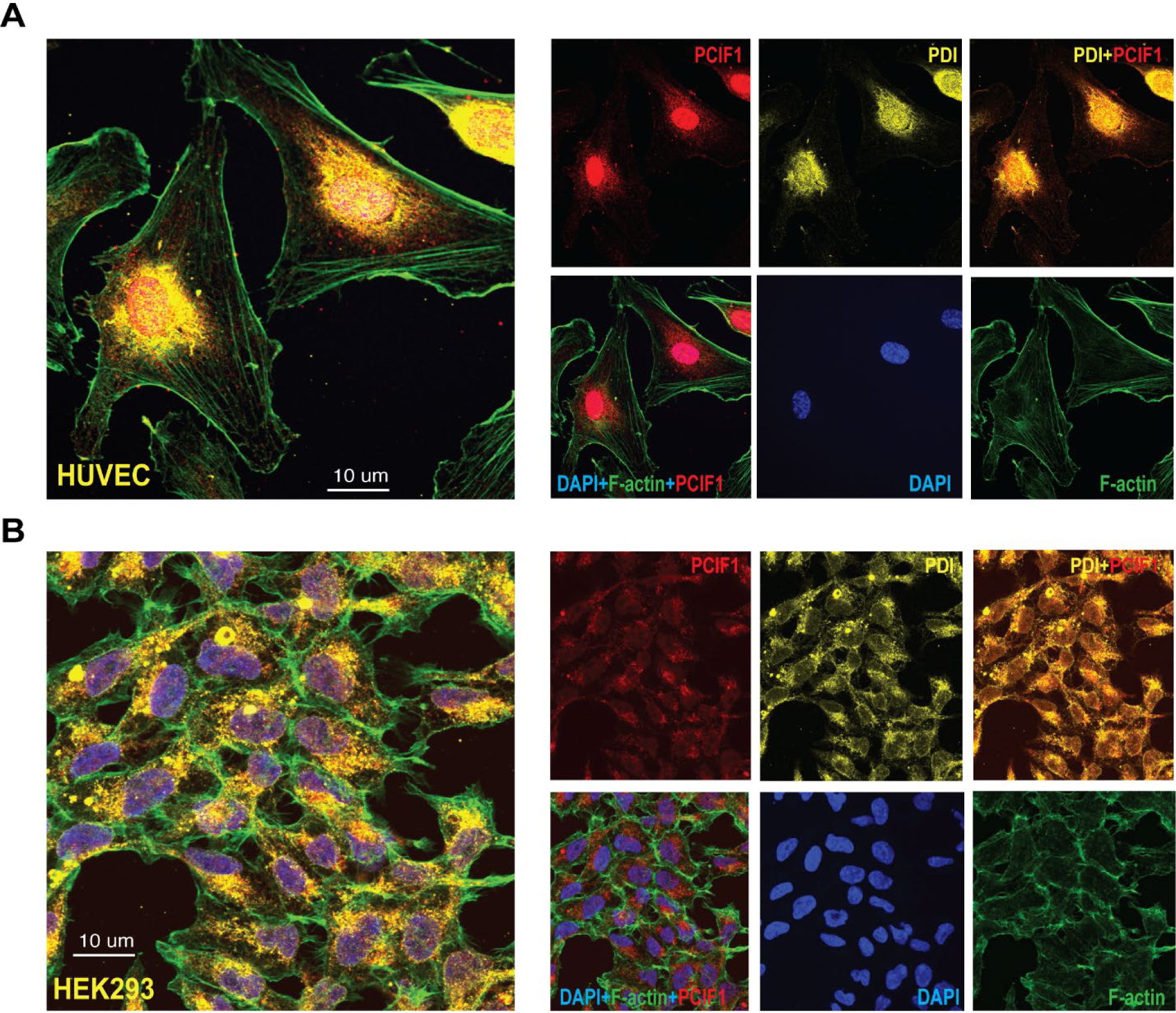
PCIF1 is found both in the cytoplasm and nuclei of two mammalian cell lines. (A) HUVEC and (B) HEK293 cells were grown on coverslips and stained with antibodies targeting PCIF1 (red), Protein disulfide isomerase (PDI, yellow), phalloidin (green) and DAPI (blue). Images represent the consensus data from three independent biological replicates.

**Supplementary Fig S4.**
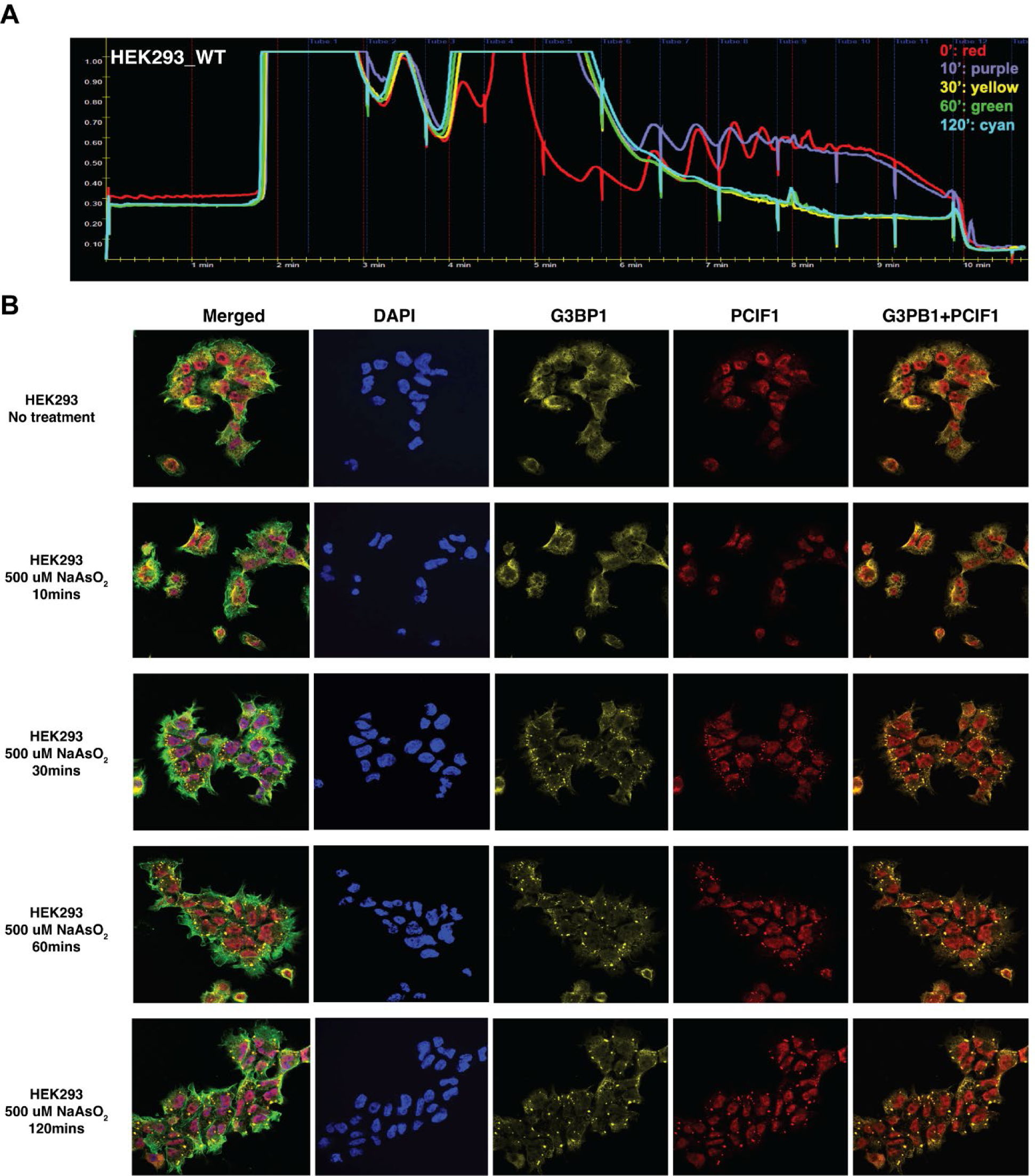
Treatment with 500 µM NaAsO_2_ induces the integrated stress response. PCIF1 is predominantly found in cytoplasmic compartment in a variety of mammalian cell types. (A) Polysome profiling was performed with 10% to 50% linear sucrose gradients as described in methods to assay global translation shutdown after HEK293 cells were treated with 500 µM NaAsO_2_ for the indicated times. Gradients were fractionated using a Brandel fractionator and A_254_ was measured continuously using the PeakChart software. (B) Indirect immunofluorescence experiments were performed as earlier and the formation of stress granules and PCIF co-localization with them is shown over time. Antibodies targeted PCIF1 (red), G3BP1 (yellow), phalloidin (green) and DAPI (blue). Images represent the consensus data from three independent biological replicates.

**Supplementary Fig S5.**
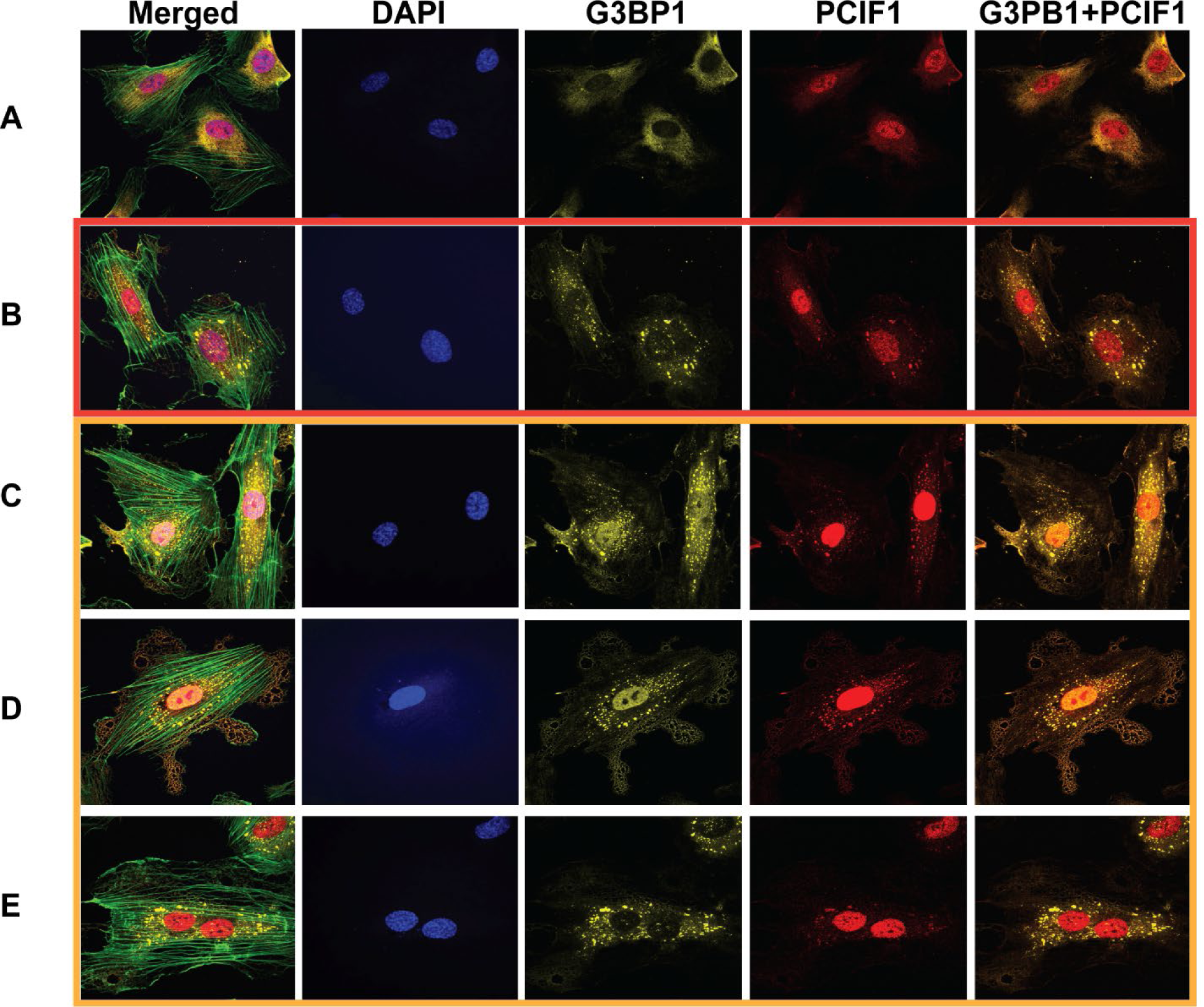
Immunofluorescent data, using 4 different commercially available PCIF1-against antibodies, revealed the presence of cytoplasmic PCIF1 in HUVEC cells. Unstressed control (A) and cells treated with 500 µM NaAsO_2_ for 60 mins (B – E) HUVEC cells were stained with antibodies targeting PCIF1 (red), G3BP1 (yellow), phalloidin (green) and DAPI (blue). The PCIF1-targeting antibody is (A & B) α-PCIF1 (Sigma, SAB1407847), (C) α-PCIF1 (Thermo Fisher, PA5-110081), (D) α-PCIF1 (Sigma, HPA049517), (E) α-PCIF1 (Santa Cruz, sc-374406). Images represent the consensus data from three independent biological replicates.

**Supplementary Fig S6.**
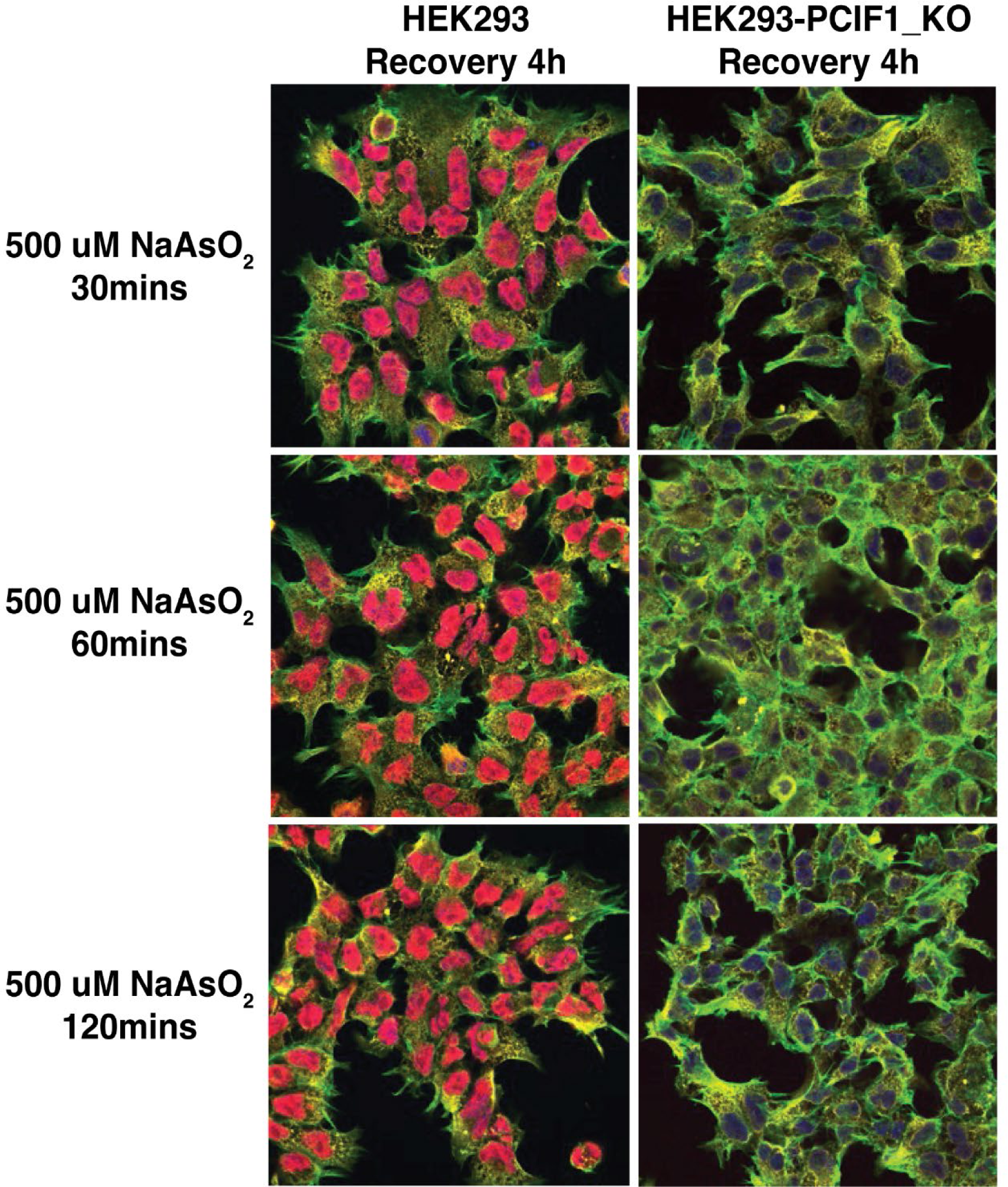
Extended stress recovery studies using WT and PCIF1^-/-^ HEK293 cells. HEK293 cells were grown on coverslips and treated with NaAsO_2_ for the indicated times. To visualize stress granule disassembly, the NaAsO_2_ was removed, and cells were allowed to recover prior to fixing and staining. Cells were fixed and stained with antibodies targeting PCIF1 (red), G3BP1 (yellow), phalloidin (green) and DAPI (blue) as described in methods. Representative merged images of each treatment showing the enrichment of PCIF1 and G3BP1 in stress granules upon NaAsO_2_ stress.

**Supplementary Fig S7.**
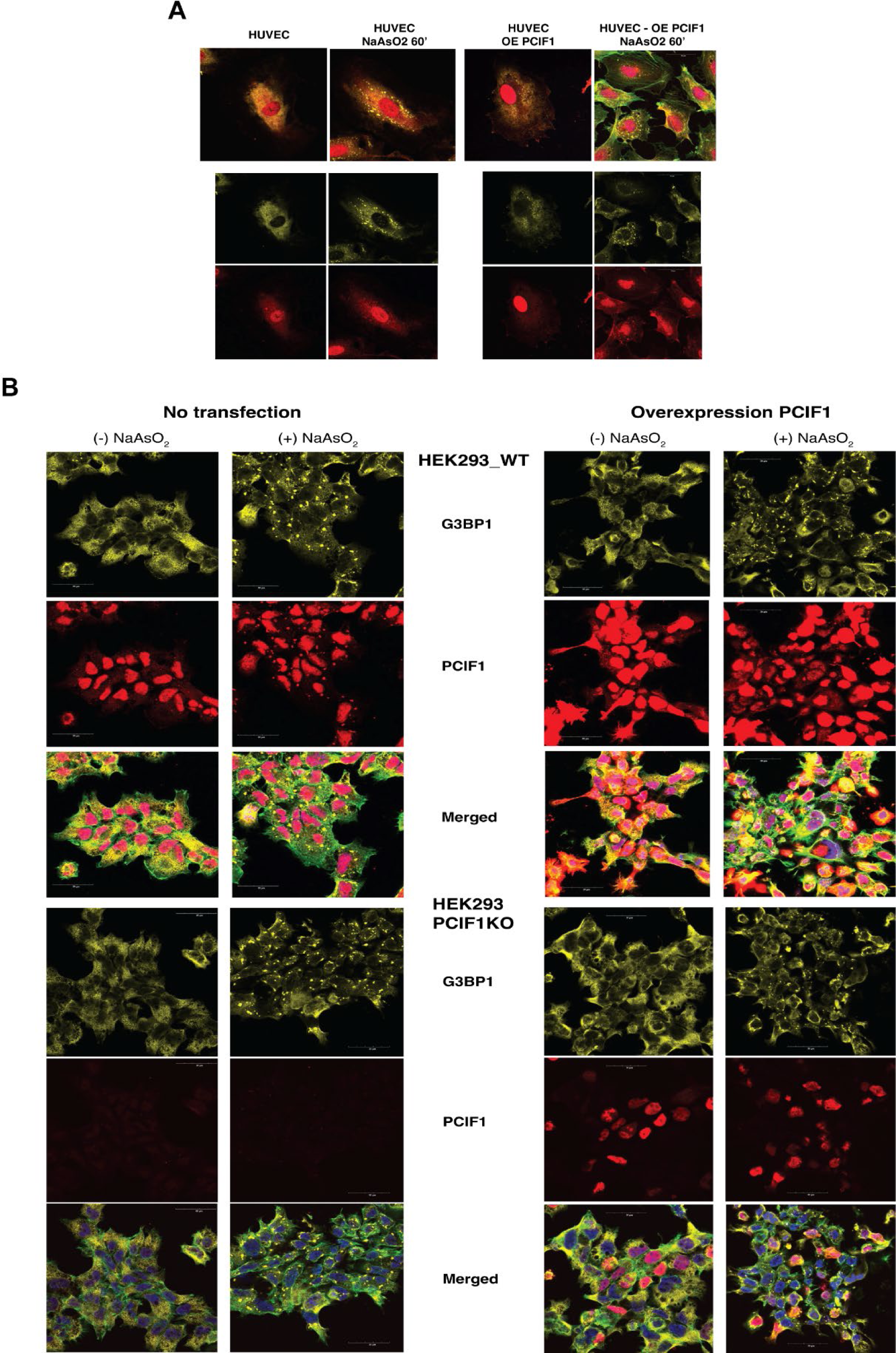
Overexpression of PCIF1 inhibits/delays stress granule formation in HUVEC and WT and PCIF1^-/-^ HEK293 cells. (A and B) HUVEC and both WT and PCIF1^-/-^ HEK293 cells were grown on coverslips and either mock transfected or transfected with a PCIF1 expression vector ∼18 hours prior to being treated with NaAsO_2_ for 60 minutes. Cells were fixed and stained with antibodies targeting PCIF1 (red), G3BP1 (yellow), phalloidin (green) and DAPI (blue) as described in methods. Transfected WT cells are characterized by intense PCIF1 staining (red). The number and size of stress granules are reduced in PCIF1 overexpressing cells. (B, bottom) Overexpressed PCIF1 is readily observable in PCIF1^-/-^ HEK293 cells. Note the decreased number of stress granules in PCIF1 overexpressing knockout cells. Images represent the consensus data from three independent biological replicates.

**Supplementary Fig S8.**
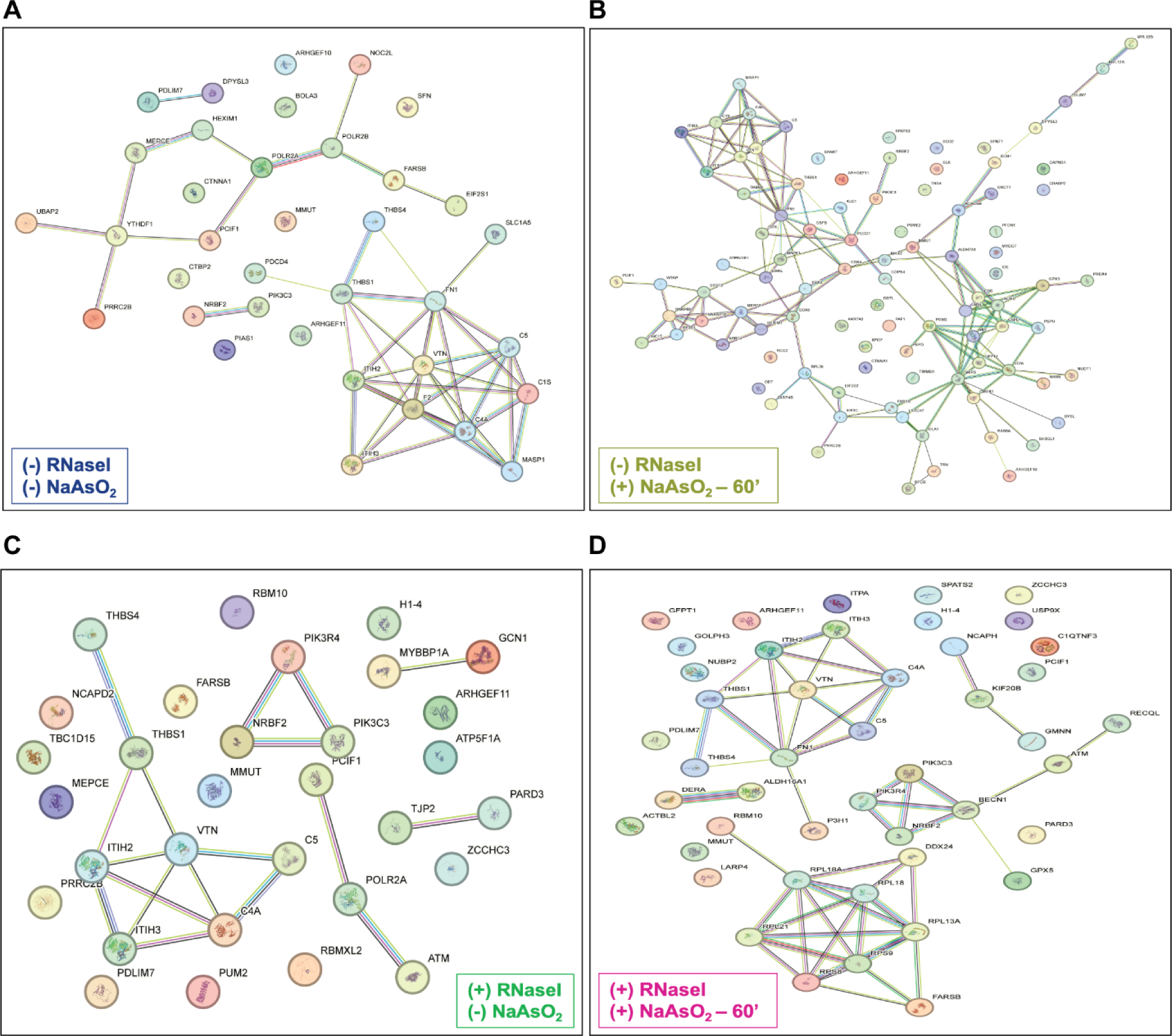
Protein interaction network maps were generated using STRING (ver 12.0) for proteins immunoprecipitated by PCIF1. The maps for all four groups (A) No RNase I, No stress, (B) No RNase I, Plus stress, (C) Plus RNase I, No stress, and (D) Plus RNase I, Plus stress, are shown.

**Supplementary Fig S9.**
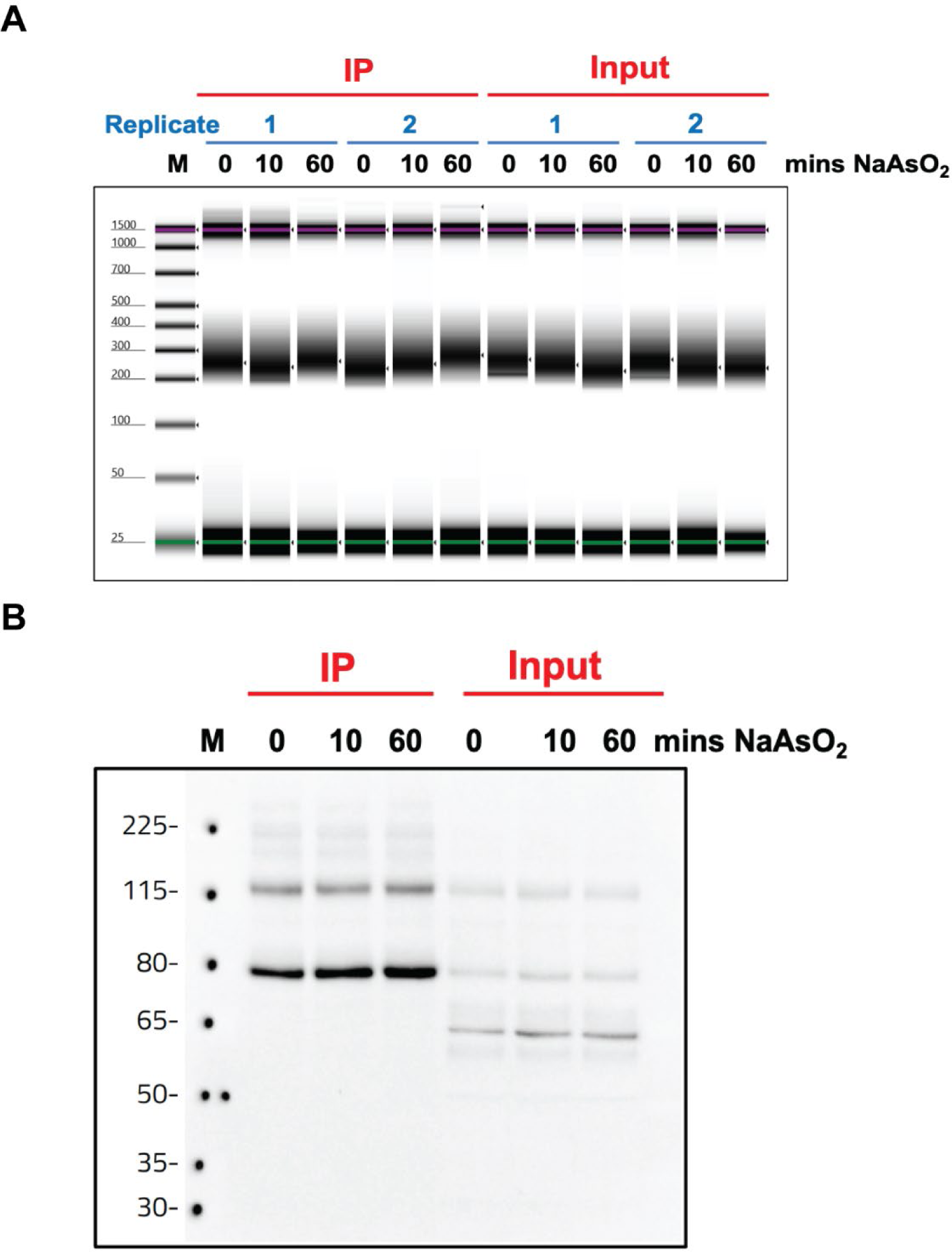
Quality control parameters for eCLIP library preparations. (A) Tapestation trace showing the sheared RNA inputs for eCLIP library preparations. (B) Western blotting of immunoprecipitation during PCIF1 eCLIP with antibody against N-PCIF1 and TrueBlot anti-Rabbit IgG (HRP). Only the region from 80 kDa to 150 kDa was isolated during eCLIP.

**Supplementary Fig S10.**
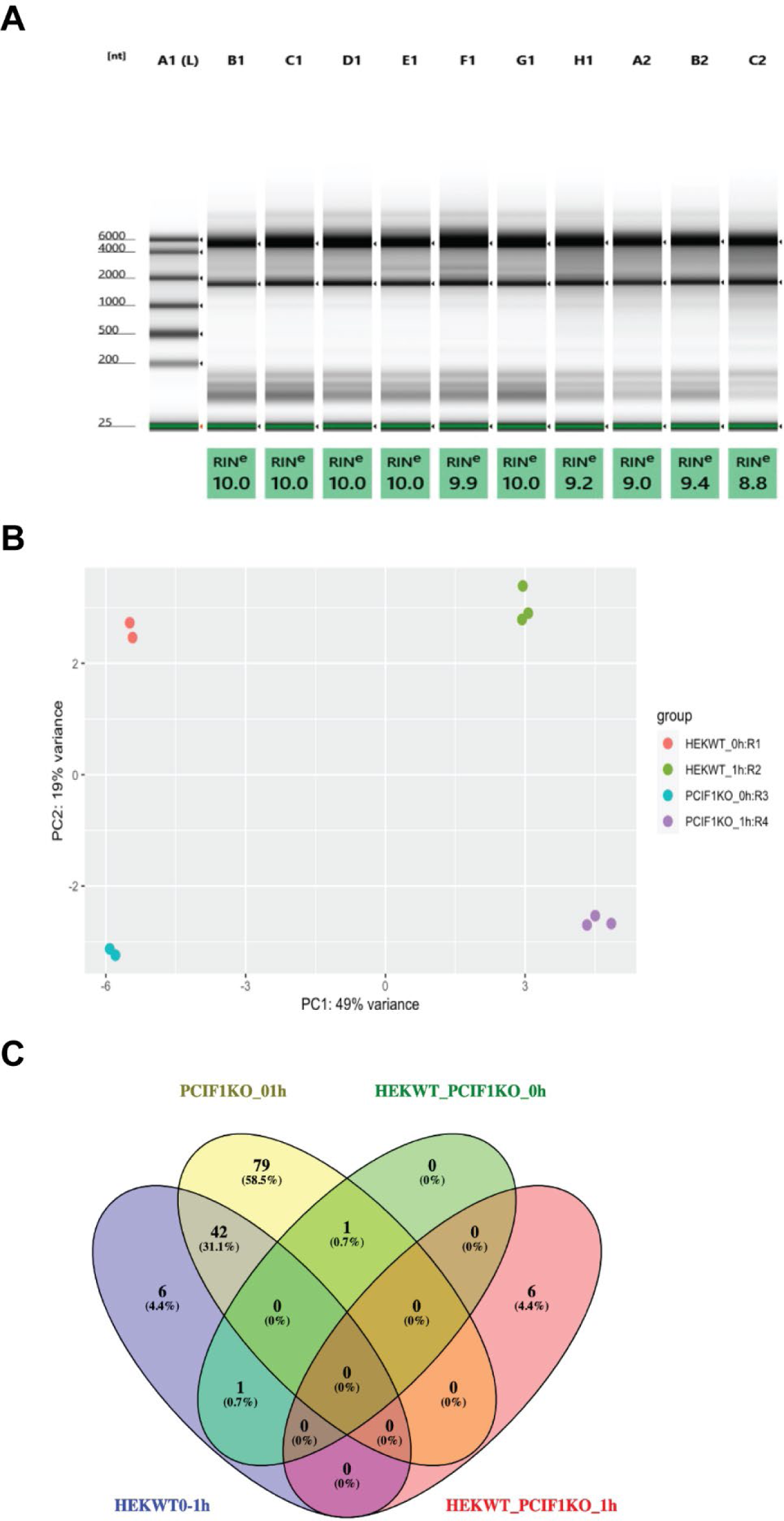
Nanopore Direct RNA Sequencing library preparations. (A) Tapestation trace showing representative samples of the total RNA inputs used in Nanopore library preparations. (B) Principal component analysis showing the consistency of independent replicates for the Nanopore direct RNA sequencing data sets. (C) A Venn diagram showing the number of shared (yet significantly changed) transcripts between the four groups of Oxford Nanopore data.

**Supplementary Fig S11.**
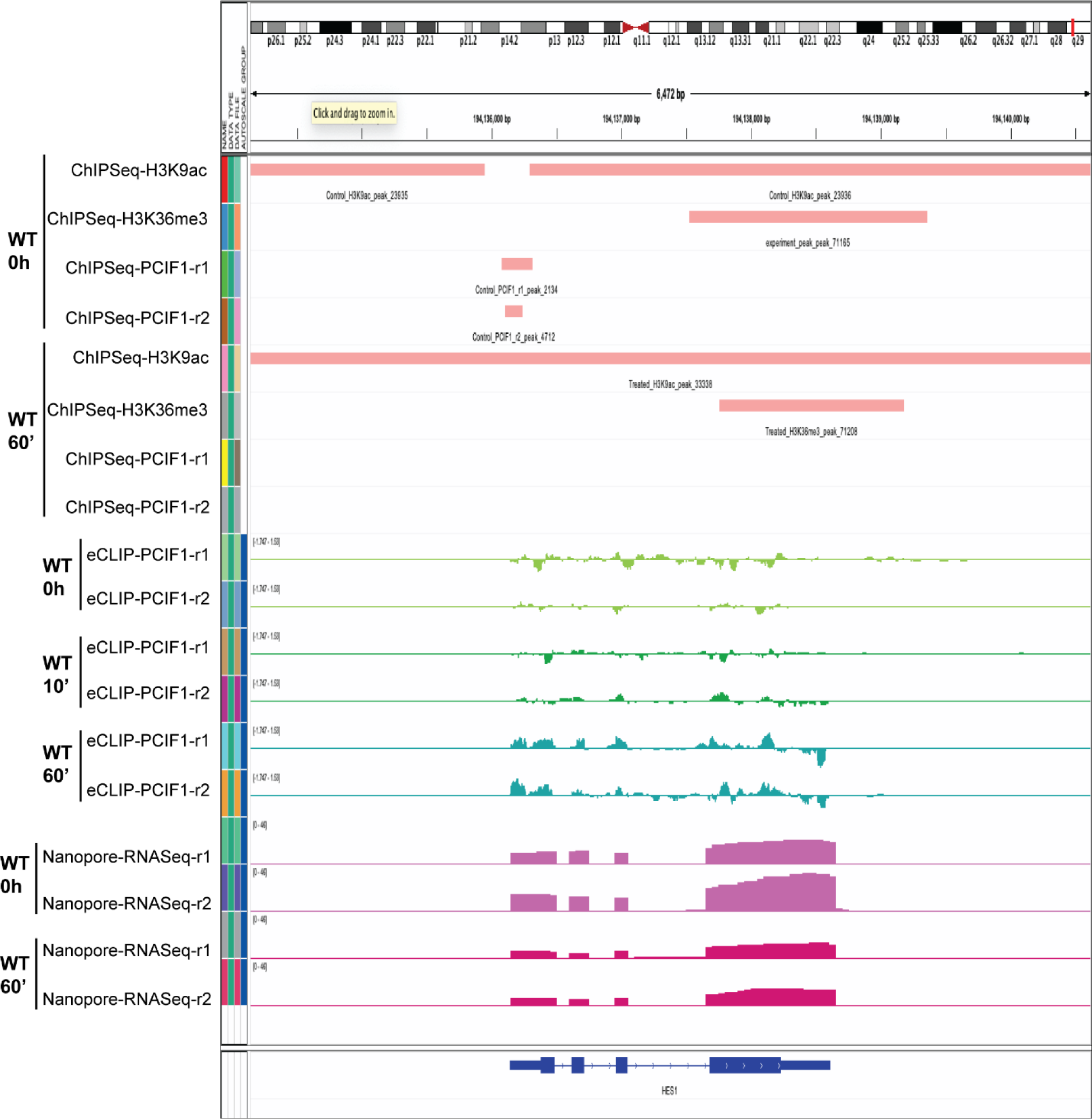
Multiple sequencing techniques show PCIF1’s association with mRNA is not co-transcriptional. Visual representation of ChIP-Seq (IP antibodies were for Histone H3K9ac, Histone H3K36me3, and PCIF1), eCLIP (with PCIF1 antibody), and Oxford Nanopore direct RNA sequencing profiles mapped to HES1 mRNA. The data shown are the consensus of ChIPSeq, eCLIP and direct RNA sequencing results from duplicate experiments. The plot was made with IGV.

**Supplementary Fig S12.**
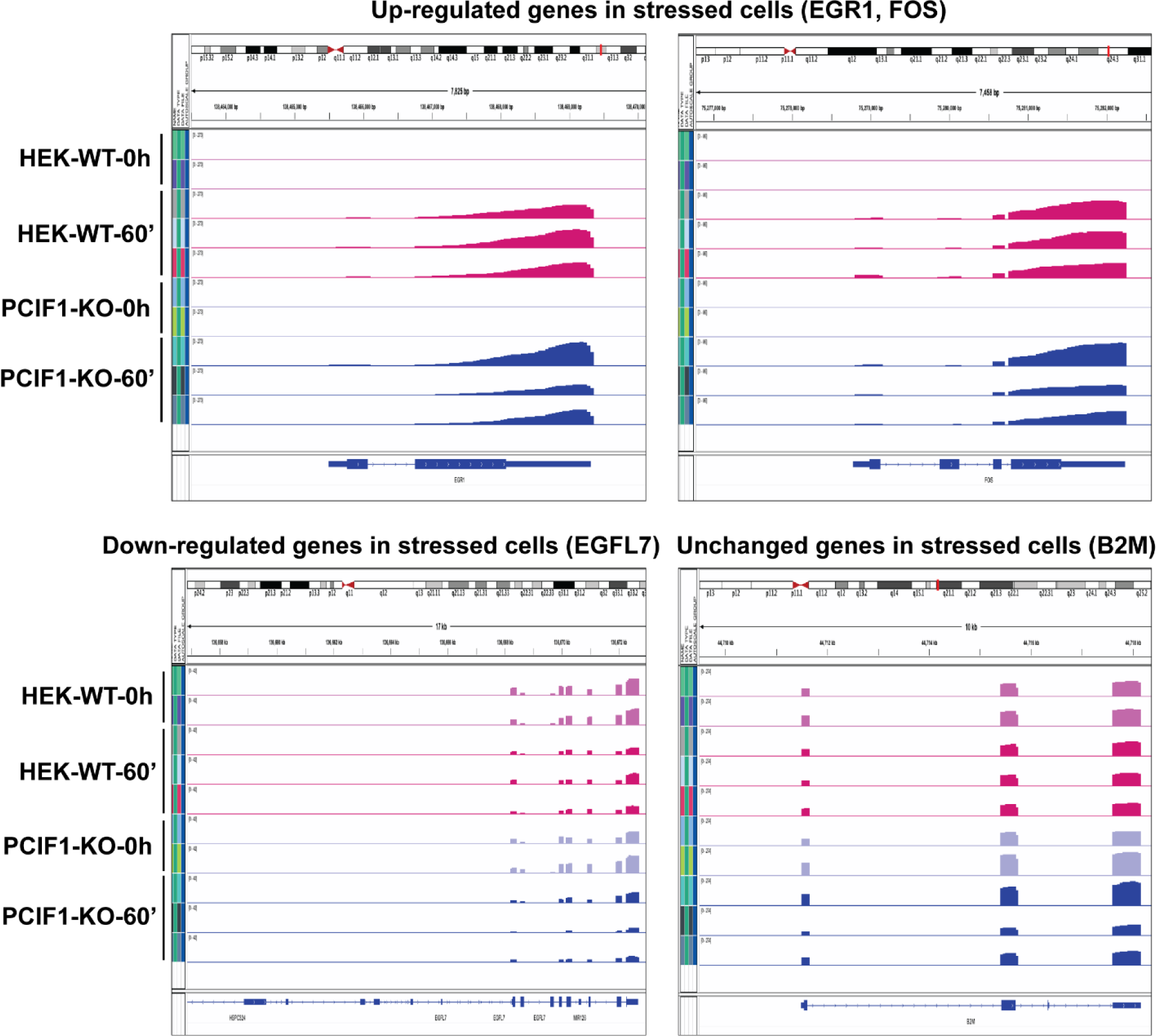
IGV visualization shows the up-, down-regulated and unchanged genes in HEK-WT and PCIF1^-/-^ HEK293 cells using Nanopore Direct RNA sequencing. The data shown are the consensus of direct RNA sequencing results from duplicate experiments.

**Supplementary Fig S13.**
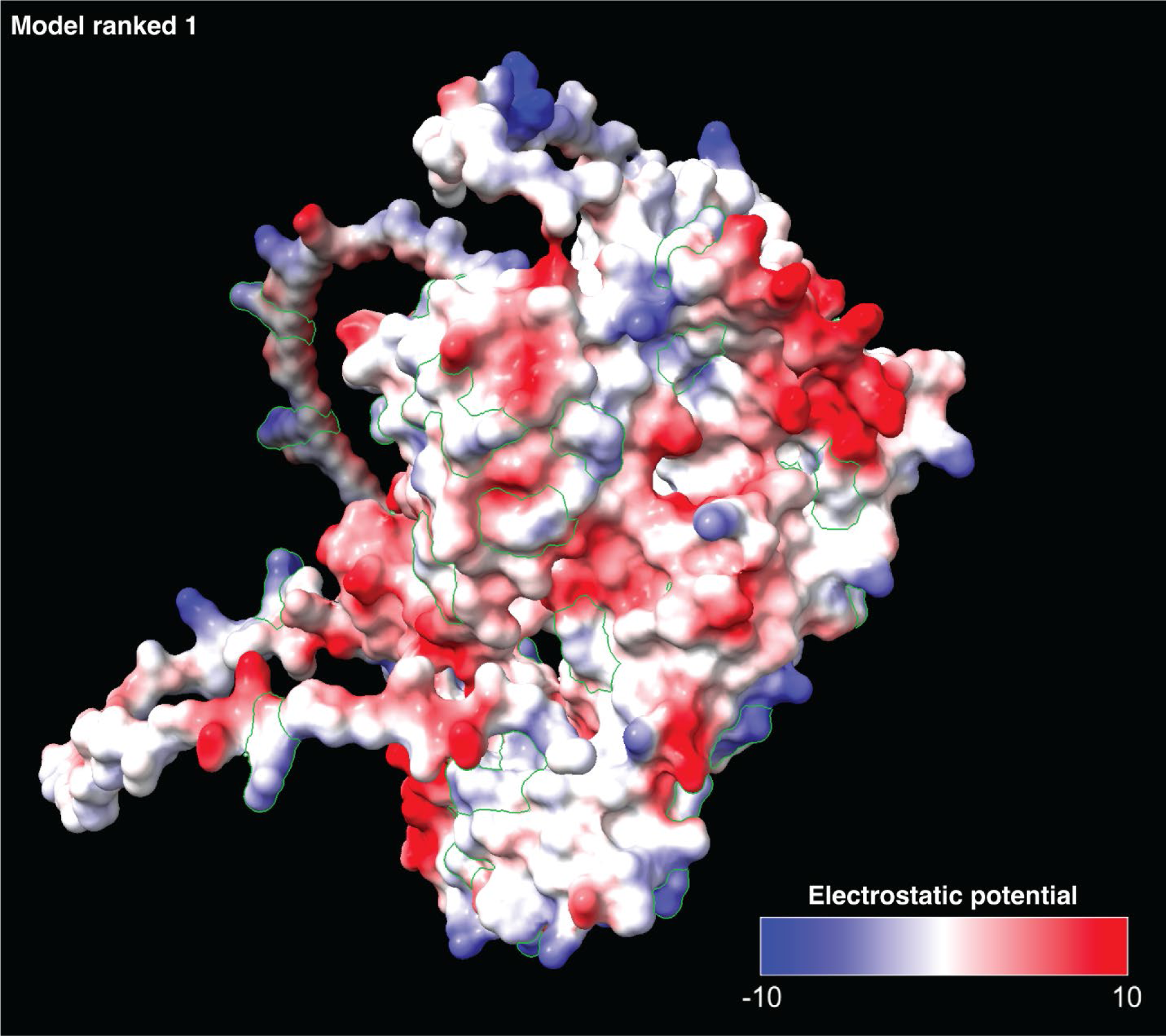
Alphafold reconstruction showing charged face of PCIF1 protein as the putative binding interface with RNA. The image was created using Alphafold ver 2 and ChimeraX-1.7.1. Positively charged Arginine residues are outlined in green.

**Supplement Table 1. Key resources table.**

**Supplement Table 2. PCIF1 Mass Spectrometry results and Comparisons to known protein components of stress granules**

**Supplement Table 3. PCIF1 eCLIP results and Comparisons to mRNAs known to localize to or be excluded from stress granules**

**Supplement Table 4. Changed mRNAs observed in WT and PCIF1^-/-^ cells as observed by Oxford Nanopore direct RNA sequencing**

