## Supplement Table 1. Key resources table for "PCIF1 is partly cytoplasmic, dynamically localizes to stress granules and binds mRNA coding regions upon oxidative stress"

**Table S1: KEY RESOURCES TABLE**

| Reagents or resources | Source | Identifier (Cat#) |
|---|---|---|
| **Antibodies** | | |
| $\alpha$-PCIF1 (against entire protein) | Sigma | SAB1407847 |
| $\alpha$-PCIF1 (against N-terminal region) | Sigma | HPA049517 |
| $\alpha$-PCIF1 (against N-terminal region, used in IF) | Thermo Fisher | PA5-61995 |
| $\alpha$-PCIF1 (against center region, used in IF) | Thermo Fisher | PA5-112227 |
| $\alpha$-PCIF1 (against N-terminal region, used in eCLIP and IP) | Thermo Fisher | PA5-110081 |
| $\alpha$-PCIF1 (against C-terminal region) | Abcam | ab205016 |
| $\alpha$-PCIF1 (against C-terminal region) | Bethyl | A304-711A |
| $\alpha$-PCIF1 (against C-terminal region) | ProteinTech | 16082-I-AP |
| $\alpha$-PCIF1 | Santa Cruz | sc-374406 |
| $\alpha$-Lamin B | Abcam | MA1-06101 |
| $\alpha$-beta Actin | ProteinTech | 66009-I-Ig |
| $\alpha$-G3BP1 | Abcam | ab56574 |
| $\alpha$-GAPDH | ProteinTech | 60004-I-Ig |
| $\alpha$-Histone H3 | Ab Clonal | A2348 |
| $\alpha$-PDI | Invitrogen | MA3-019 |
| $\alpha$-PDI | Cell Signaling | C81H6 |
| $\alpha$-IgG Rabbit (used in IP) | Milipore | 12-370 |
| Goat AntiMouse IgG (H+L) Alexa Fluor Plus 647 | Thermo Fisher | A32728 |
| Goat AntiRabbit IgG (H+L) Alexa Fluor Plus 647 | Thermo Fisher | A32733 |
| Goat AntiMouse IgG (H+L) Alexa Fluor Plus 594 | Thermo Fisher | A11032 |
| Goat AntiMouse IgG (H+L) Alexa Fluor Plus 555 | Thermo Fisher | A32727 |
| Goat AntiRabbit IgG (H+L) Alexa Fluor Plus 555 | Thermo Fisher | A32732 |
| HPR-conjugated $2^{nd}$ antibody | Cell Signaling | 7074S |
| $2^{nd}$ anti-Mouse True Blot | Rockland | 18-8817-33 |
| $2^{nd}$ anti-Rabbit True Blot | Rockland | 18-8816-33 |
| $\alpha$-H3K36me3 | Abcam | ab9050 |
| $\alpha$-H3K9ac | Diagenode | C15410004 |
| **Bacterial strains** | | |
| GC5 DH5$\alpha$ | NEB | C2987U |
| **Reagents** | | |
| EBM$^{TM}$ Basal Medium | Lonza | CC-3121 |
| EGM$^{TM}$ Endothelial Cell Growth Medium SingleQuots Supplements | Lonza | CC-4133 |
| 100X Pen/Strep | Gibco | 15140-122 |
| DMEM (high glucose) | ATCC | 30-2002 |
| FBS | Corning | 35-075-CV |

| | | |
|---|---|---|
| McCoy's 5A medium | Gibco | 16600-082 |
| Zeocin | Invitrogen | 46-0509 |
| Blasticidin S HCl 10 mg/ml | Gibco | A11139-03 |
| L-glutamin | Gibco | 25030-081 |
| OptiMEM I (1X) Reduced Serum Media | Gibco | 11058-021 |
| Trypsin-EDTA 0.25% | Gibco | 25200-056 |
| DMSO | Sigma | D4540 |
| Lipofectamin 3000 | Invitrogen | L3000001 |
| JetMessenger mRNA Transfection Reagent | Polyplus | 150-07 |
| GlycoBlue Coprecipitant | Invitrogen | AM9515 |
| Dynabeads M280 sheep antiRabbit IgG | Thermo Fisher | 11204D |
| Dynabeads M280 sheep antiMouse IgG | Thermo Fisher | 11201D |
| Protein Inhibitor Cocktail (PIC) | Sigma | P8340-5ml |
| Phosphatase Inhibitor Cocktail II | Sigma | P5726 |
| Phosphatase Inhibitor Cocktail III | Sigma | P0044 |
| PMSF 0.1M | Sigma | 93482-50ml-F |
| NP40 10% | Thermo Fisher | 28324 |
| Phosphate Buffered Saline (PBS) | GenClone | 25-507B |
| F-actin Alexa Fluor 488 Phallodin | Thermo Fisher | A12379 |
| Diamond antifade mountant with DAPI | Thermo Fisher | P36966 |
| Fibronectin coated German Glass Cover Slips | Electron Microcopy Sciences | 72297-05 |
| MycoStrip™ – Mycoplasma Dectection Kit | InvivoGen | Rep-mysnc-50 |
| 4-15% Mini-PROTEAN TGX Precast Protein Gels | Biorad | 4568086 |
| Biorad Protein Assay Dye Reagent Concentrate | Biorad | 5000006 |
| 10X Tris/Glycine/SDS Buffer | Biorad | 1610732 |
| TransBlot Turbo 5X Transfer buffer | Biorad | 10026938 |
| PVDF membrane | Thermo Fisher | 22860 |
| Tris Buffer Saline, Tween-20 (TBST 10X) pH7.6 | Apex | 18-235B |
| Instant Nonfat Dry Milk | RPI | M17200-500.0 |
| Ethanol 200 Proof | Decon Labs, Inc | 64-17-5 |
| Precision Plus Protein Dual Color Standards | Biorad | 1610374 |
| MPER Solution | Thermo Fisher | 78505 |
| Restore Plus Western Blot Stripping Buffer | Thermo Fisher | 46430 |
| $NaAsO_2$ | Santa Cruz | sc-301816 |
| Paraformaldehyde (PFA) 16% Solution, EM Grade | Electron Microcopy Sciences | 15710 |
| BSA | Fisher | BP9703-100 |
| Triton X-100 | Sigma | T8787 |
| Emetine | Calbiochem | 324693 |
| $MgCl_2$ | Invitrogen | AM9530G |
| Sucrose | Sigma | S9378 |
| TRI Reagent | Zymo Research | R2050-1-200 |
| Chloroform | Sigma | C2432 |
| DTT 1X | Sigma | 646563 |
| RNase-In Plus | Promega | N261B |
| Clarity™ Western ECL Substrate | Bio-rad | 1705061 |
| Acid acetic, glacial | VWR | 0714-500ml |
| GridION flow cells | Oxford Nanopore Technologies | R9.4.1 |
| **Critical commercial assays** | | |

| | | |
|---|---|---|
| Plasmid maxiprep kit | Zymo Research | D4203 |
| Dynabeads Co-Immunoprecipitation kit | Thermo Fisher | 14321D |
| Pierce Silver Stain Mass Spectrometry | Thermo Fisher | 24600 |
| IP Antibody Validation Kit | Eclipse BioInnovations | ECAV-0001 |
| Direct-zol RNA Miniprep | Zymo Research | R2052 |
| Monarch RNA Cleanup Kit (500 ug) | NEB | T0250L |
| Nanopore Direct RNA Sequencing Kit | Nanopore | SQK-RNA002 |
| **Experimental models** | | |
| **Cell lines** | | |
| Primary Human Umbilical Vein Endothelial Cells (HUVEC) | Lonza | #CC-2517 |
| Flp-In T-Rex HEK293 | Gibco | R78007 |
| Flp-In T-Rex HEK293 – PCIF1 KO | Synthego | 9582438-1 |
| U2OS | | |
| **Recombinant DNA** | | |
| PCIF1 (NM_022104) Human Tagged ORF Clone | OriGene | RC203430 |
| **Software and algorithms** | | |
| AlphaFold v.2.3.2 | | |
| bedGraphToBigWig v.366 | | |
| Bedtools v.2.30.0 | | |
| CLIPper v.1.3.0 (https://github.com/YeoLab/clipper) | | |
| Cutadapt v.3.2 | | |
| DESeq2 v.1.43.5 | | |
| FastQC v.0.11.9 | | |
| FV31S-SW v2.6 | | |
| gpGrouper algorithm v.1.0 | | |
| HTSeq v. 0.12.4 | | |
| IGV v.2.17.4 | | |
| ImageLab v.4.1 | | |
| MACS2 v.2.1.2 | | |
| Mascot ver.2.4 | | |
| Matplotlib v.3.3.4 | | |
| Minimap2 v.2.28 | | |
| MinKNOW v.1.4.2 | | |
| NanopPot v.1.41.0 | | |
| Numpy v.1.19.2 | | |
| Pandas v.1.1.5 | | |
| PD v.2.1 | | |
| Prism v.7.04 | | |
| Proteome Discoverer v.2.1 | | |
| Pybedtools v.0.8.1 | | |
| Pysam v.0.15.0.1 | | |
| Python v.3.6.6 :: Anaconda v.4.10.1 (64-bit) | | |
| R v. 4.4.0 | | |
| R::Bioconductor v. 3.19 | | |
| SAMtools v.1.9 | | |
| STAR v.2.7.7a | | |
| Statistics::Basic v.1.6611 | | |
| Statistics::Distributions v.1.02 | | |
| Statistics::R v.4.4.0 | | |
| UCSF ChimeraX v.1.7.1 | | |

| | |
|---|---|
| Human RefSeq database (release Jan. 21, 2020) | |
| **Equipments** | |
| Eppendorf table top microfuge | Eppendorf |
| Bioruptor$^{R}$ Pico | Diagenode |
| ChemiDoc MP imaging System | Bio-rad |
| Olympus FV3000 Confocal Microscopy | Olympus |
| Brandel gradient fractionator | Brandel |
| UV Crosslinker | UVP, CL-1000 |
| TapeStation 4200 | Agilent Technologies |
| Illumina NextSeq 500 | Illumina |
| Nanopore GridION X5 | Oxford Nanopore Technologies |
| Orbitrap Fusion mass spectrometers | Thermo Fisher Scientific |
| Easy-nLC 1000 nanoflow LC system | Thermo Fisher Scientific |
